## Supplementary Materials for "Efficient and sustained optogenetic control of sensory and cardiac systems"

### Supplementary Material

List of supplementary material:

1. Supplementary Figure 1: Peak recovery of ChRmine variants.
2. Supplementary Figure 2. Automated photocurrent measurements.
3. Supplementary Figure 3: Noise analysis of ChReef and ChRmine photocurrents.
4. Supplementary Figure 4: Voltage dependence of closing kinetics of ChRmine variants.
5. Supplementary Figure 5: Subcellular localization of ChRmine and ChReef in NG cells
6. Supplementary Figure 6: Action spectra of ChRmine wt and ChReef.
7. Supplementary Figure 7: Assessment of relative permeabilities of ChRmine and ChReef.
8. Supplementary Figure 8: Dependence of photocurrents of NG cells expressing ChRmine and ChReef on the intensity of light of different colors.
9. Supplementary Figure 9: Comparison of photocurrent densities of HEK293T cells expressing ChReef and ChRmine in response to red light of different intensities.
10. Supplementary Figure 10 Optical pacing and depolarization block of hIPS-derived CM clusters.
11. Supplementary Figure 11: ChReef expression in other brain regions in animals for vision restoration experiments
12. Supplementary Figure 12: Characterization of oABR P1-N1 amplitudes among stimulation with 594nm.
13. Supplementary Figure 13: Immunohistochemical analysis of ChReef expression in the AAV-injected gerbil cochlea.
14. Supplementary Figure 14: Inferior colliculus recordings in ChReef-injected Mongolian gerbils.
15. Supplementary Figure 15: Temporal resolution of optogenetic activation of the auditory pathway by means of inferior colliculus recordings in Mongolian gerbils.
16. Supplementary Figure 16: Comparison of oABR from Figure 5 and published aABR data.
17. Supplementary Figure 17: Quantification of hair cell loss following deafening of the injected left side in the common marmoset.
18. Supplementary Table 1. Stationary-Peak-Ratio, $EC_{50}$ values, stationary current densities and $\tau_{off}$ values of ChRmine variants.
19. Supplementary Table 2. Stationary current densities and $\tau_{off}$ values of ChR variants.
20. Supplementary Table 3. Exact (Mann-Whitney U-test) and adjusted (Bonferroni) p-values from the Figure 1a.
21. Supplementary Table 4. Exact (Mann-Whitney U-test) and adjusted (Bonferroni) p-values from the Figure 1b.
22. Supplementary Table 5. Exact (Student's t-test) and adjusted (Bonferroni) p-values from the Figure 1c.
23. Supplementary Table 6. Exact p-values (Tukey's HSD test) from the Figure 1i.
24. Supplementary Table 7. List of primers used for ChRmine mutant generation.
25. Supplementary Table 8. Solutions used for acute slice in vitro electrophysiology.
26. Supplementary Table 9. Exact (Mann-Whitney U-test) and adjusted (Bonferroni) p-values from the Figure 5b.
27. Supplementary Table 10. Exact (Mann-Whitney U-test) and adjusted (Bonferroni) p-values from the Figure 5c.

28. Supplementary Table 11. Exact p-values (Tukey's HSD test) from the Supplementary Figure 5d.

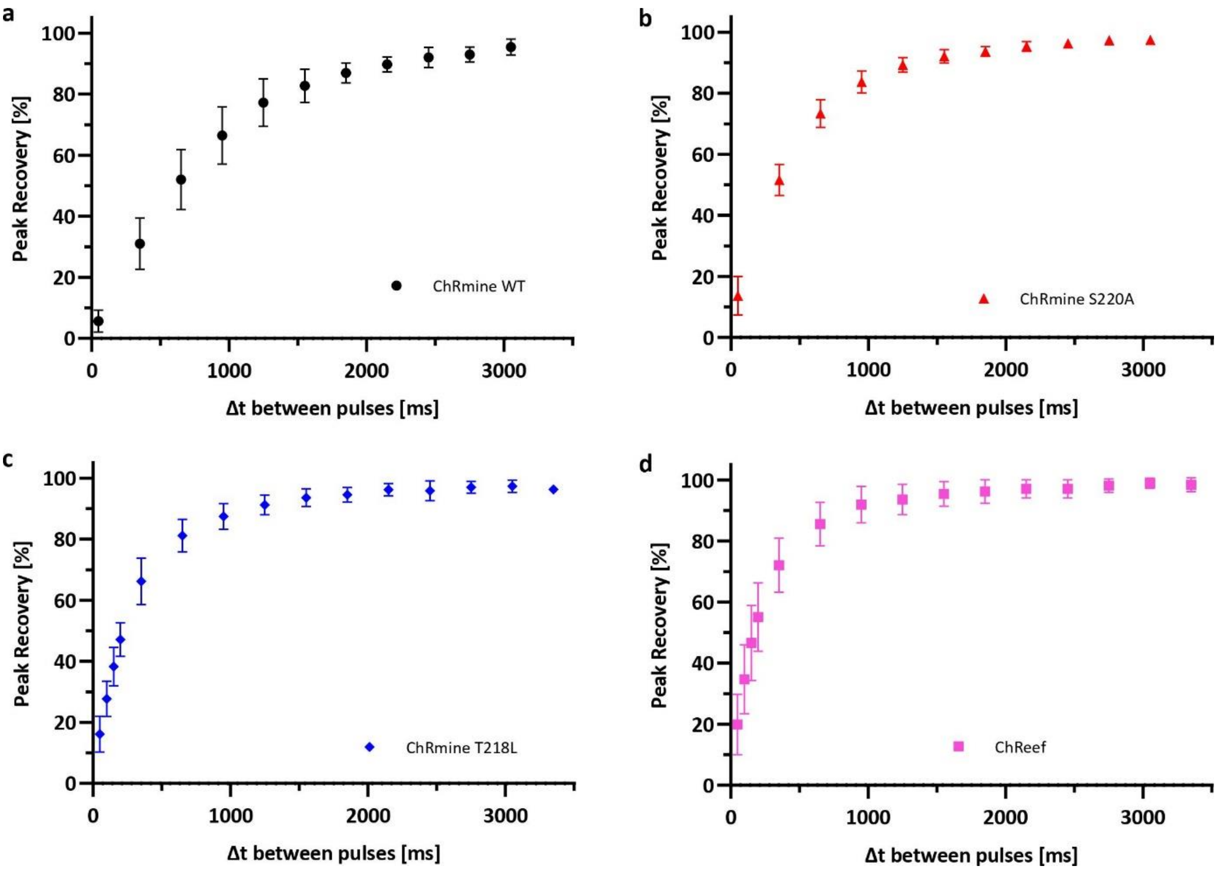


**Supplementary Figure 1: Peak recovery of ChRmine variants.** NG cells transiently transfected with **(a)** ChRmine WT (black circle, n=4), **(b)** ChRmine S220A (red triangle, n=4), **(c)** ChRmine T218L (blue rhombus, n=5) and **(d)** ChReef (magenta square, n=5) were investigated by whole-cell patch-clamp recordings at a membrane potential of -60 mV. Photocurrents were measured upon illumination with two subsequent 3 s light pulses of a wavelength of $\lambda$=532 nm at a light intensity of 23 mW/mm$^2$ with a varying time between the light pulses (Δt between pulses, ms). Peak recovery was calculated as the quotient of the difference between the peak and stationary current of pulse two ($I_{p2}$-$I_{s2}$) and peak and stationary current of pulse one ($I_{p1}$-$I_{s1}$) in %. The symbols indicate the mean, error bars show SD, n indicates the number of measured cells.

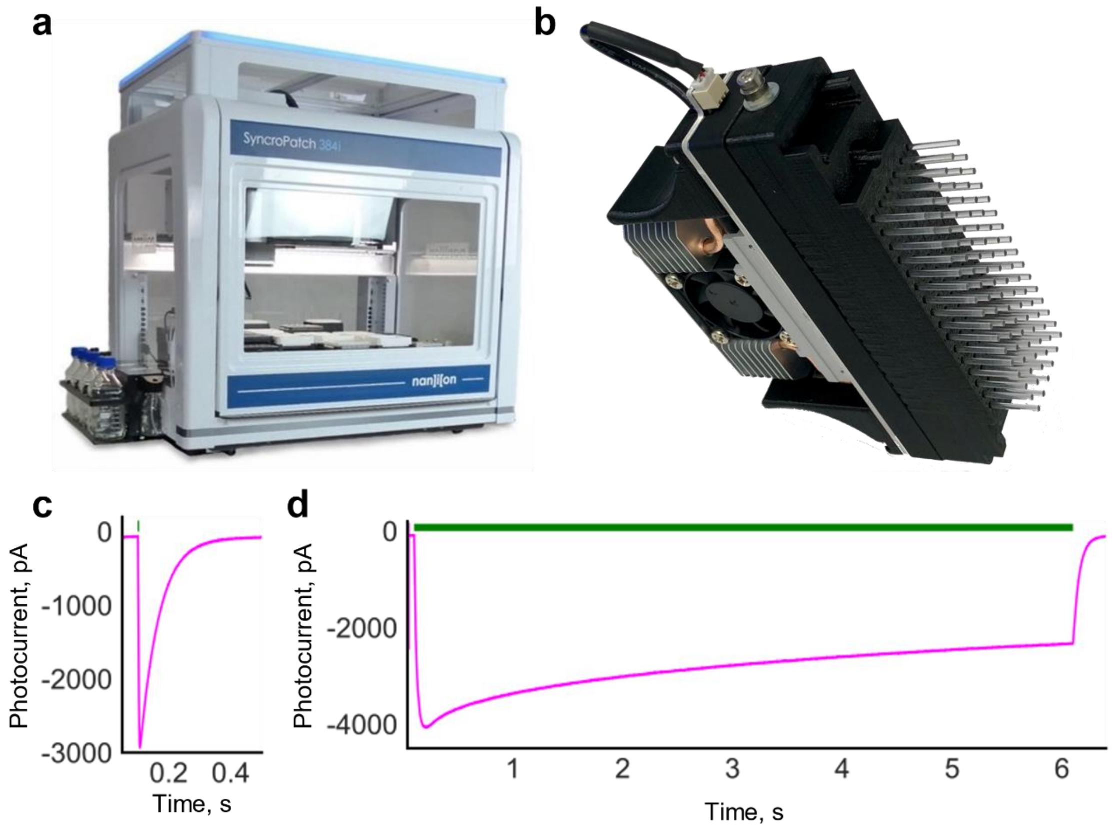


**Supplementary Figure 2. Automated photocurrent measurements.** **a**, Picture showing the high-performance automated patch-clamp system SyncroPatch384 (Nanion) **b**, Picture showing the illumination unit (Nanion), which comprises 96 LEDs coupled to 96 individual light fibers and a cooling system with a fan. **c**, Exemplary ChReef photocurrent at a membrane potential of -100 mV evoked by a 5-ms green light pulse, which was used for non-stationary noise analysis. **d**, Exemplary ChReef photocurrent at a membrane potential of -100 mV evoked by a 6-s green light pulse, which was used for stationary noise analysis.

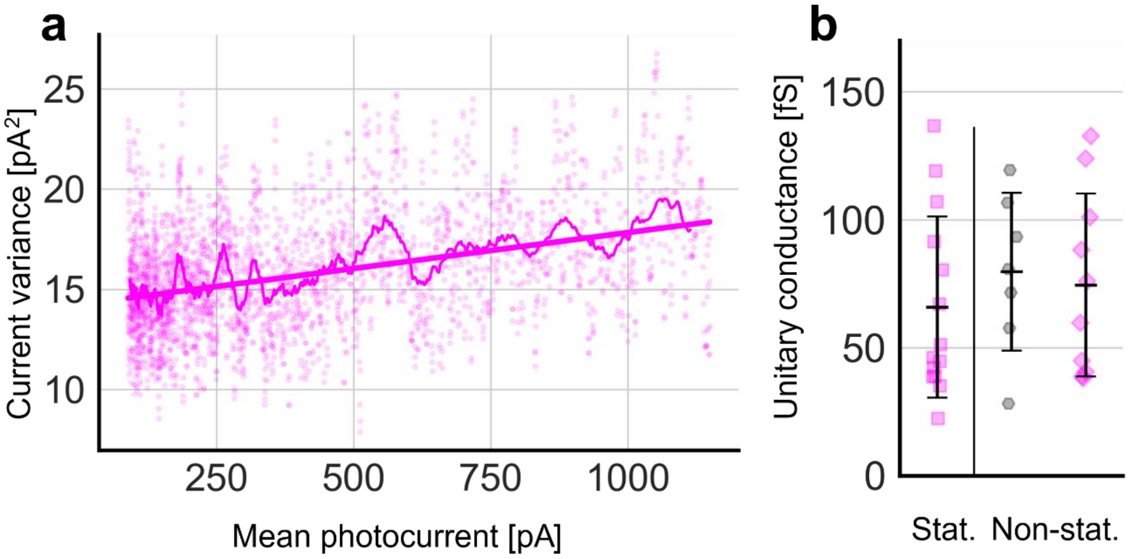


**Supplementary Figure 3. Noise analysis of ChReef and ChRmine photocurrents a**, Non-stationary noise analysis of ChReef. Exemplary plot showing the relation between the variance of the photocurrent and its average value. The photocurrents were measured in automated whole-cell patch-clamp experiments at a membrane potential of -60 mV. The thin line shows the moving average of the variance. The bold line indicates a linear fit. **b**, Statistical comparison of single channel conductance values at a membrane potential of -60 mV. The single channel conductance values of ChReef (magenta diamonds, n=10) and ChRmine (grey hexagons, n=7) derived from non-stationary noise analysis are shown on the right. On the left side the single channel conductance value of ChReef (magenta squares, n=14) derived from stationary noise analysis is shown. n indicates the number of measured cells.

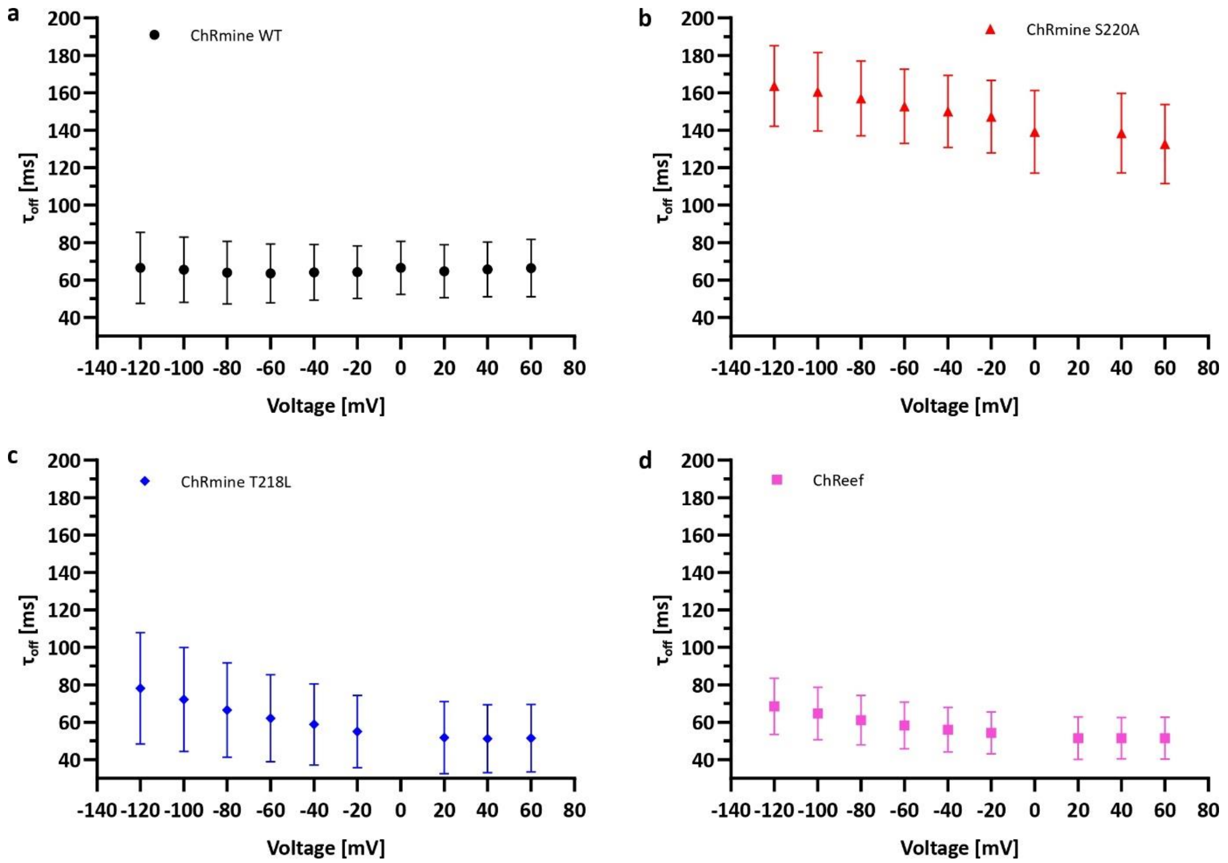


**Supplementary Figure 4: Voltage dependence of closing kinetics of ChRmine variants.** NG cells transiently transfected with ChRmine (black circle, WT), ChRmine S220A (red triangle), ChRmine T218L (blue rhombus) and ChReef (magenta square) were investigated by whole-cell patch-clamp recordings at different membrane potentials ranging from -120 mV to 60 mV. Photocurrents were measured upon illumination with a 3 ms light pulse of a wavelength of $\lambda$=532 nm at a light intensity of 23 mW/mm$^2$. The closing kinetics ($\tau_{off}$ value) were determined by a fit of the decaying photocurrent elicited in response to the light pulse to a monoexponential function. Shown are the time constants of the closing kinetics $\tau_{off}$ for **(a)** ChRmine WT (n=7), **(b)** ChRmine S220A (n=6), **(c)** ChRmine T218L (n=7) and **(d)** ChReef (n=7). The symbols indicate the mean, error bars show SD, n indicates the number of measured cells.

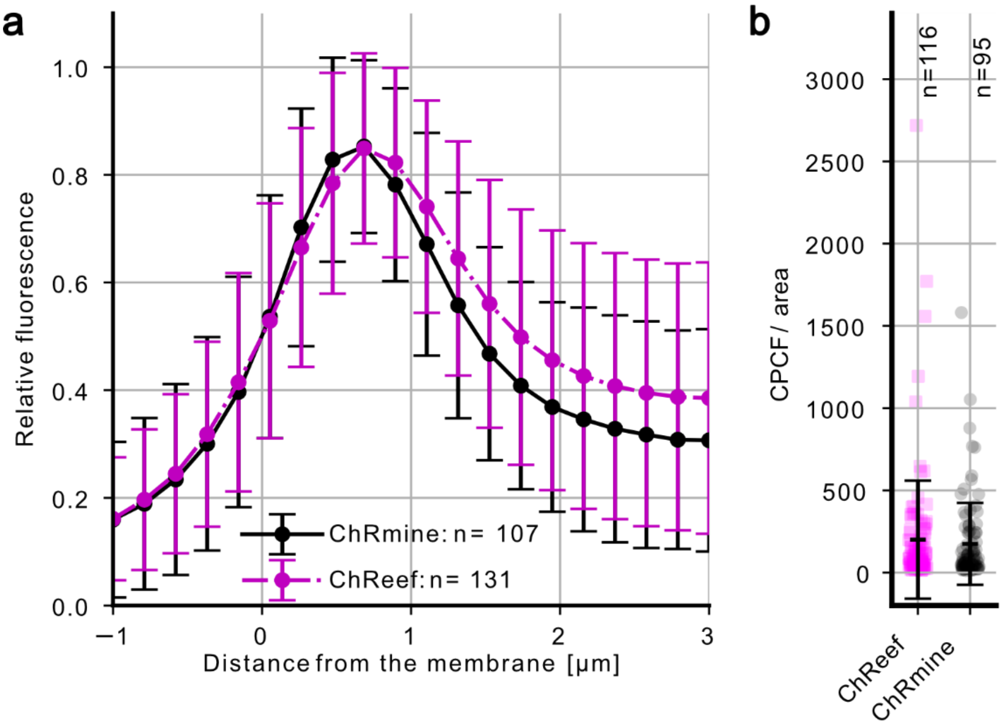


**Supplementary Figure 5: Subcellular localization of ChRmine and ChReef in NG cells.** NG cells transiently transfected with ChRmine-TS-EYFP-ES (black circle, WT) and ChReef-TS-EYFP-ES (magenta square) were imaged by automated spinning-disc confocal microscopy (Yokogawa CQ1). **(a)** Fluorescence line profile analysis of ChRmine-TS-EYFP-ES and ChReef-TS-EYFP-ES in NG cells. Shown is the average relative EYFP fluorescence intensity as a function of the estimated distance from the plasma membrane. Markers represent mean values, and vertical bars indicate standard deviation (SD). **(b)** Quantification of total protein abundance, represented by corrected partial cell fluorescence per area (CPCF/area), for NG cells expressing ChReef-TS-EYFP-ES and ChRmine-TS-EYFP-ES. Horizontal and vertical bars indicate mean values and standard deviation (SD), n indicates the number of measured cells.

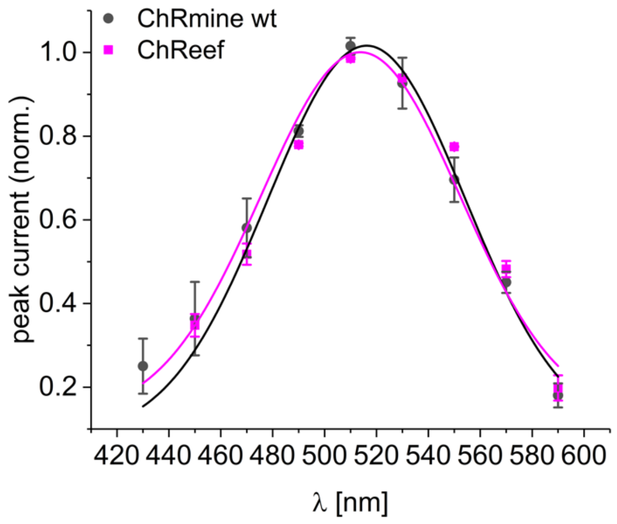


**Supplementary Figure 6: Action spectra of ChRmine wt (black circles, n=3) and ChReef (magenta squares, n=3).** Shown are normed peak currents in response to ns light-pulses of the indicated wavelengths measured in whole-cell patch-clamp experiments at a membrane potential of -60 mV. The pulse energies at the different wavelengths were set to equal photon counts. The symbols indicate average values, the error bars show SD, n indicates the number of measured cells.

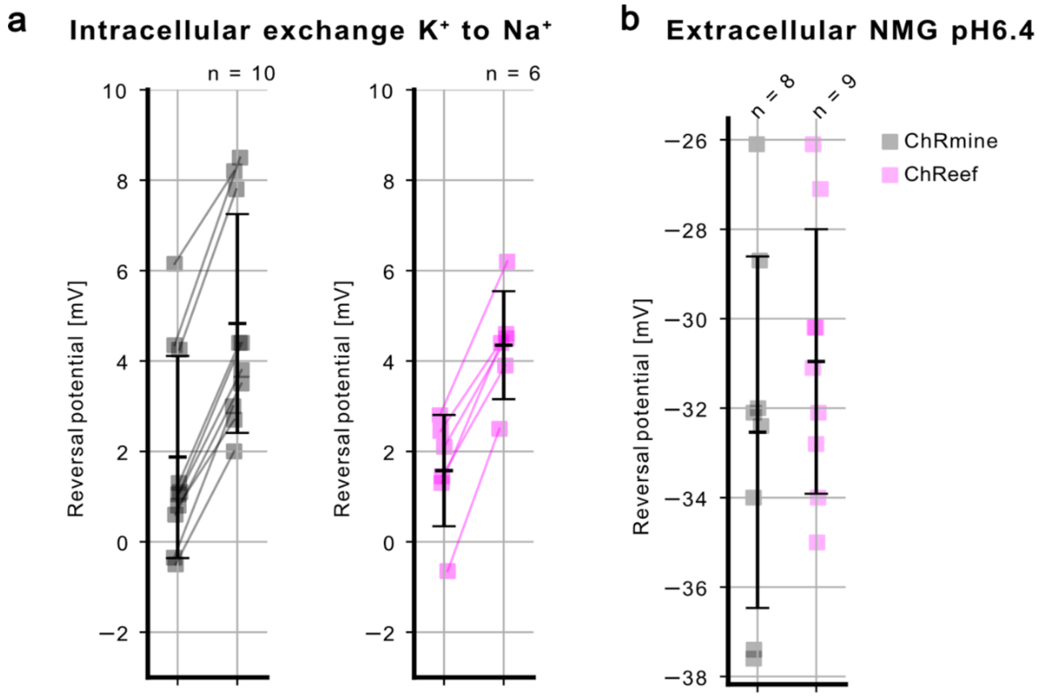


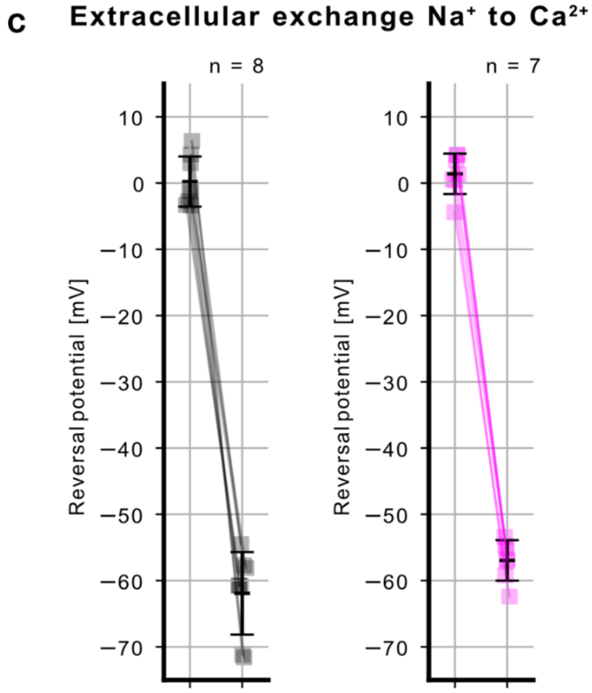


d

| | ChRmine | ChReef |
|---|---|---|
| $P_K/P_{Na}$ | 1.15±0.04 | 1.14±0.03 |
| $P_H/P_{Na}$ | ~$1.2 \cdot 10^5$ | ~$1.2 \cdot 10^5$ |
| $P_{Ca}/P_{Na}$ | n.d. | n.d. |

**Supplementary Figure 7: Assessment of relative permeabilities of ChRmine and ChReef.** **(a)** Reversal potential shifts as a result of intracellular solution exchange from high $K^+$ concentration to high $Na^+$ for estimation of relative $K^+$ permeability. **(b)** Reversal potentials of photocurrents of NG cells measured in standard high $K^+$ intracellular solution (pH 7.2) and NMG-based extracellular solution with increased proton concetration (pH 6.4) **(c)** Reversal potential shifts due to extracellular solution exchange from high $Na^+$ concentration to high $Ca^+$ for estimation of relative $Ca^+$ permeability. **(d)** Summary table with permeabilities of different cations relative to $Na^+$ permeability. n indicates the number of measured cells.

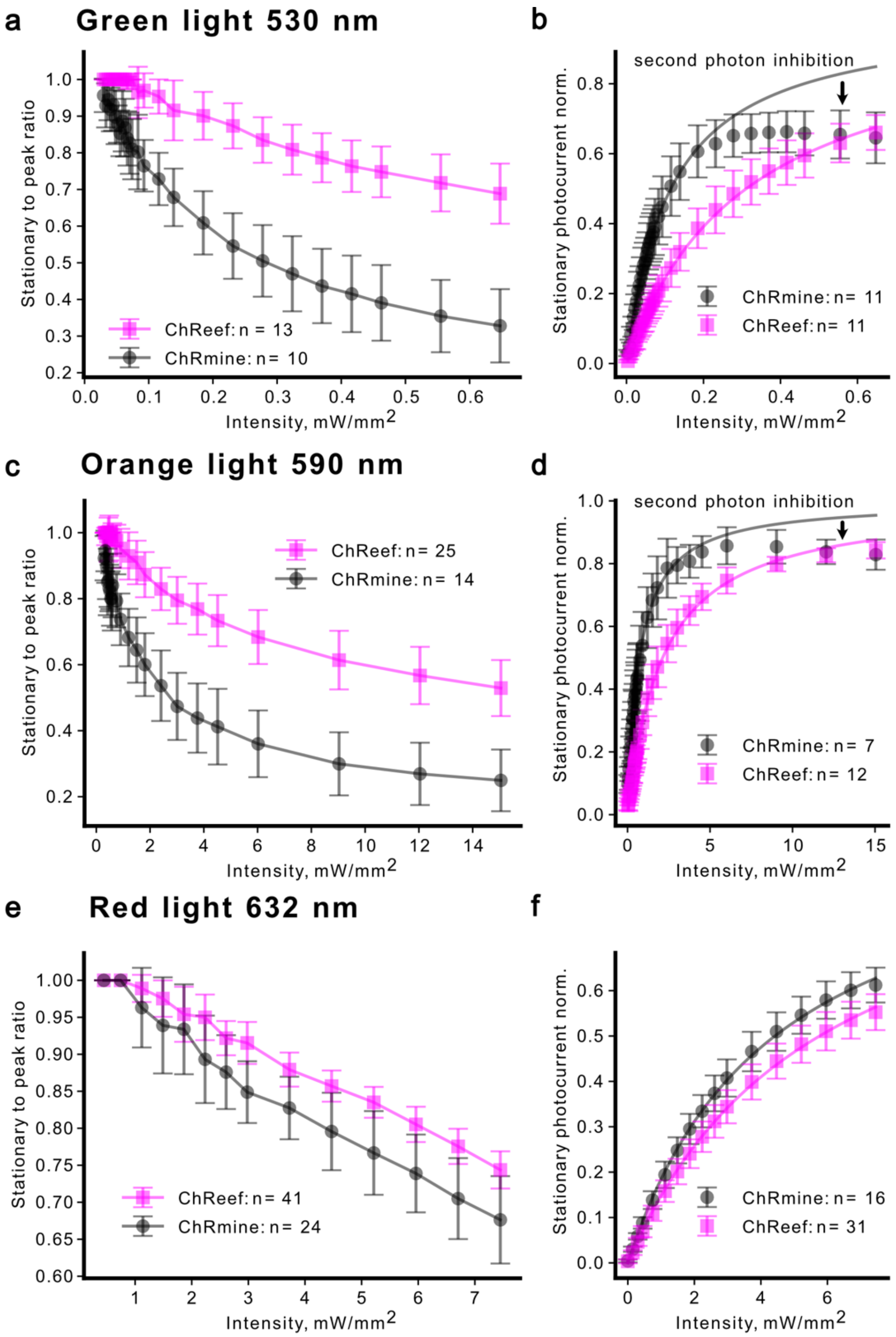


**Supplementary Figure 8: Dependence of photocurrents of NG cells expressing ChRmine and ChReef on the intensity of light of different colors. (a, b)** Green light (LED 530 nm) intensity dependence of the photocurrent peak-to-stationary ratios **(a)** and stationary photocurrent sizes **(b)**. **(c, d)** Orange light (LED 590 nm) intensity dependence of the photocurrent peak-to-stationary ratios **(c)** and stationary photocurrent sizes **(d)**. **(e, f)** Red light (LED 632 nm) intensity dependence of the photocurrent peak-to-stationary ratios **(e)** and stationary photocurrent sizes **(f)**. Markers represent mean values and error bars represent SD in all panels. n indicates the number of measured cells.

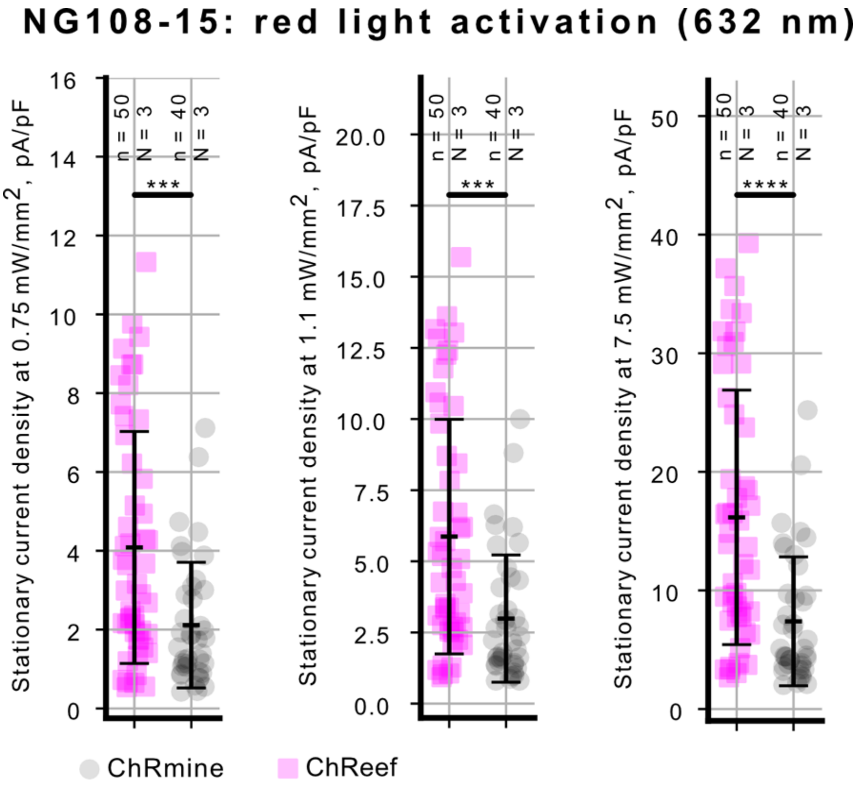


**Supplementary Figure 9. Comparison of photocurrent densities of HEK293T cells expressing ChReef and ChRmine in response to red light of different intensities.** Numbers at the top represent number of individual cells (small n) and number of transfections on different days (big N). The statistical comparisons were done with Mann-Whitney U-test (p-value < 0.05 (*), 0.005 (**), 0.001 (***), 0.0001(****)).

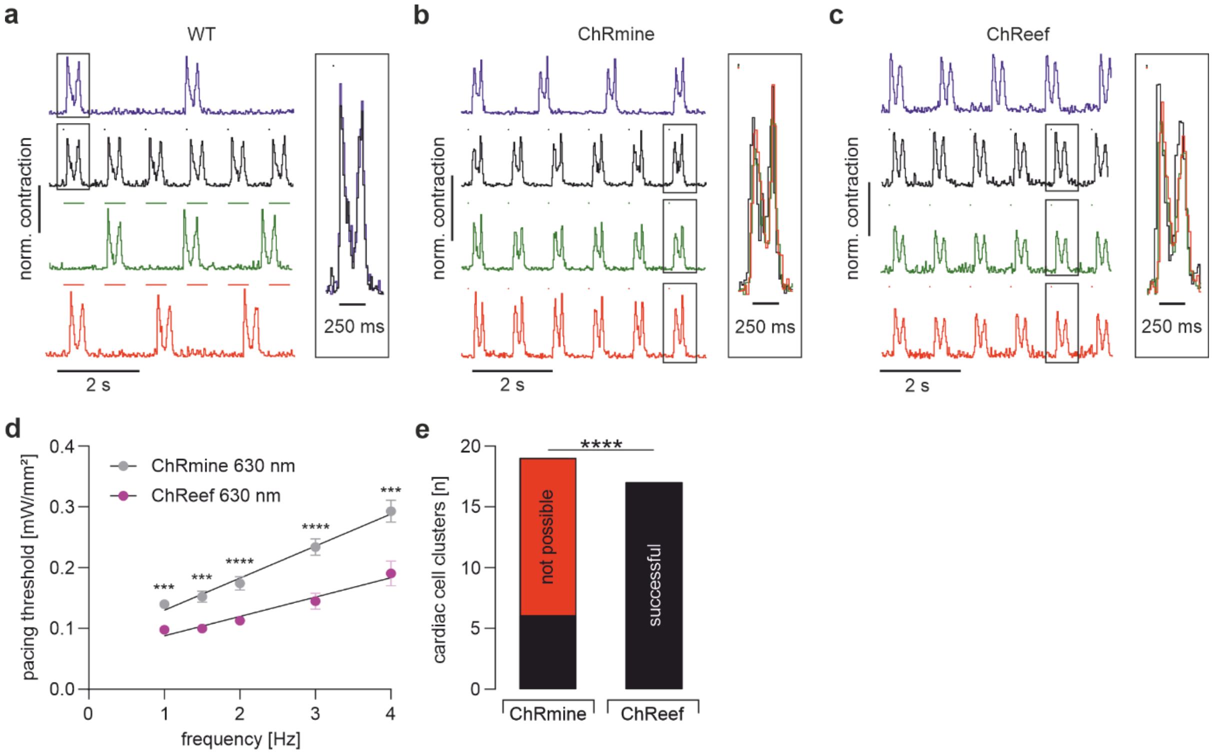


**Supplementary Figure 10. Optical pacing and depolarization block of hIPSC-derived CM clusters**

**a-c**, Representative contractions of wild type (a, for 36 CM clusters), ChRmine (**b**) and ChReef (c) expressing CM cluster (representative for n numbers indicated in d and e) during spontaneous beating (blue), upon electrical stimulation (black; dots: 0.2 ms biphasic, 25 V), green light illumination (green; dots: 510 nm, 1 ms, 1.9 mW/mm² (**a**); 8.5 µW/mm² (**b, c**) ) and red light illumination (red; dots: 630 nm, 1 ms, 5.2 mW/mm² (**a**); , 0.5 mW/mm² (**b**); 0.4 mW/mm² (**c**)). **d**, Relationship between optical pacing threshold (630 nm) and pacing frequency for ChRmine (black, n= 19) and ChReef (pink, n = 17)). Statistical comparison between ChRmine and ChReef with an unpaired t-test and significances displayed for the individual frequencies (all p values < 0.001). Analysis of the slope factor reveals a significantly steeper relationship for ChRmine (two-sided student's t-test p = 0.04). **e**, Aggregated data of the efficiency in preventing contractions with red light (maximal light intensity: 5.2 mW/mm²) for ChRmine and ChReef expressing CM clusters. Statistical analysis performed with Fisher´s exact contingency test (p < 0.0001; N = 3 MyoAAV transductions; n = 19 (ChRmine expressing CM clusters) n = 17 (ChReef expressing CM clusters)). *** indicate p-value < 0.001 and **** indicate p-value < 0.0001;

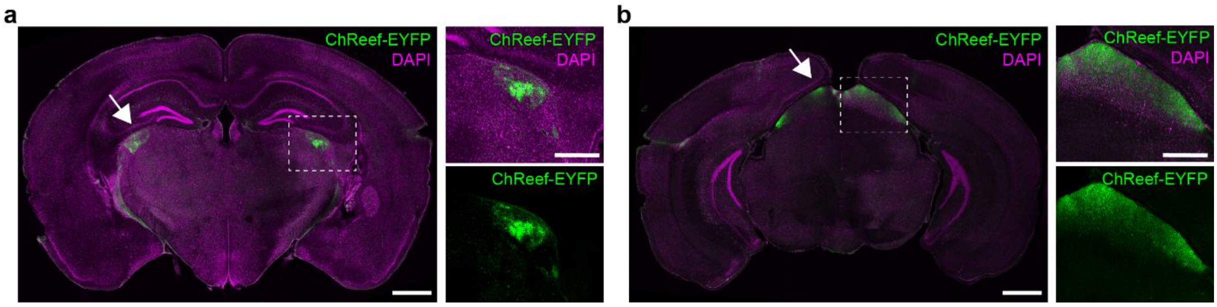


**Supplementary Figure 11. ChReef expression in other brain regions in animals for vision restoration experiments**

**a-b** Example coronal brain slices from an animal with ChReef injection in the retina. We find no expression of ChReef in the brain except for RGC axons in the lateral geniculate nucleus (rectangle and arrow in **a**) and superior colliculus (rectangle and arrow in **b**). Scale bars represent 1mm (whole slice) and 0.5mm (zoom in).

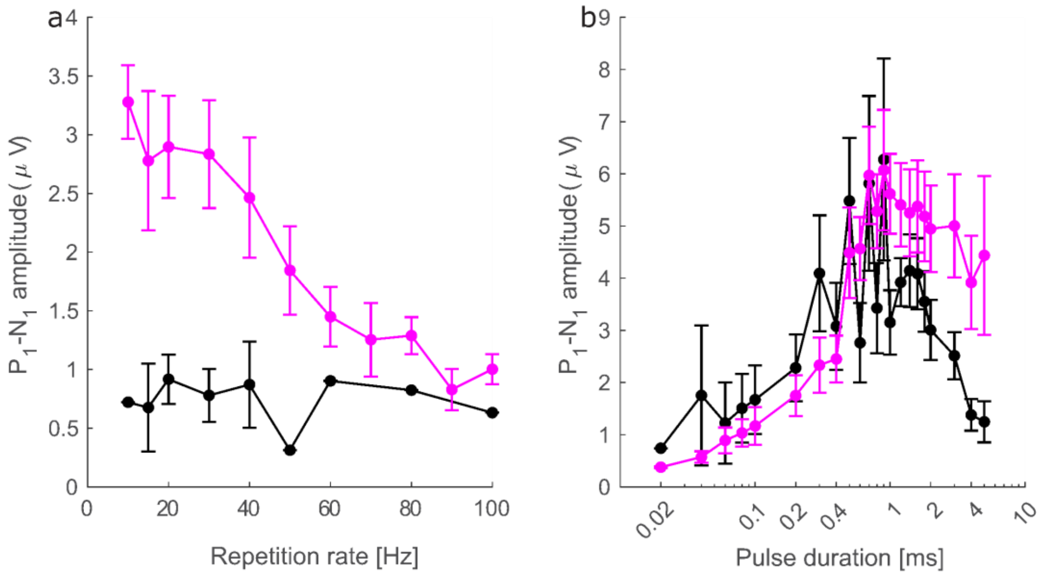


**Supplementary Figure 12. oABR P1-N1 amplitudes for different rates of stimulation with 594 nm laser light**

**a**, P1-N1 amplitudes with increasing repetition rates (for 1 ms pulses at ~10 mW radiant flux) in mice expressing ChRmine (black, N=12) or ChReef (magenta, N=15) in SGNs. **b,** P1-N1 amplitudes for varying pulse durations (at 10 Hz and ~10 mW radiant flux) in mice expressing ChRmine (black, N=12) or ChReef (magenta, N=15) in SGNs.

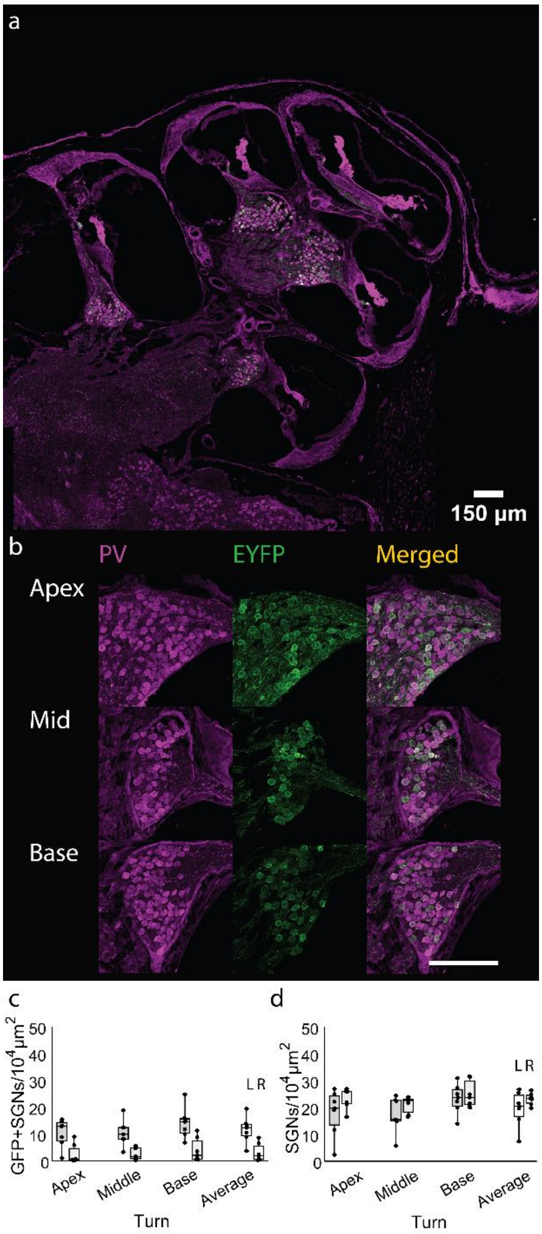


**Supplementary Figure 13. Immunohistochemical analysis of ChReef expression in the AAV-injected gerbil cochlea.**

**a**, Overview of exemplary cryosection of an AAV-injected left cochlea imaged at 20x displays expression in spiral ganglion neurons (SGNs) across all turns (representative of N=7 ChReef-injected cochleae) **b**, Exemplary cryosection of an AAV-injected left cochlea imaged at 40x shows ChR expression in SGNs across all cochlear turns (representative of N=7 ChReef-injected cochleae, staining: anti-parvalbumin (PV) as context marker, anti-GFP detecting ChR-EYFP and merged shown from left to right resp.) **c**, Boxplot computing the density of ChR-expressing SGN cells (GFP+) to the area of Rosenthal's canal observed on slice (left and right cochleae as indicated, L: injected left cochleae, N=7 gerbils, R: right non-injected cochleae, N=7 gerbils). Cell counting was performed using a custom MATLAB tool that performs assisted automatic selection of SGN. SGNs are delineated and ChR expressing GFP+ SGNs contrasted to non-expressing SGNs through principal component analysis. **d,** Boxplot computing total SGN density (left and right cochleae as indicated, L: injected left cochleae, N=7 gerbils, R: right non-injected cochleae, N=7 gerbils).

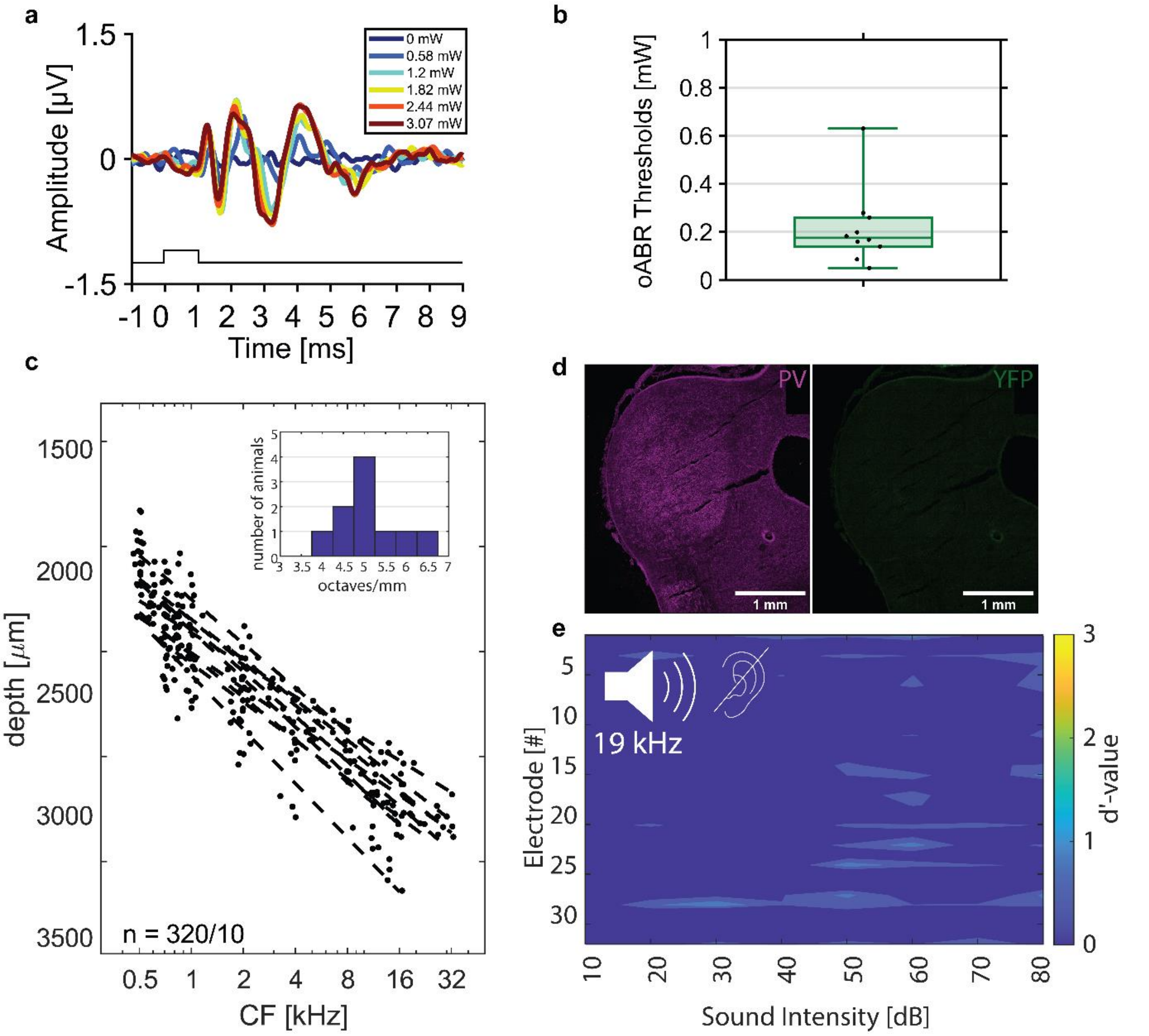


**Supplementary Figure 14. Inferior colliculus recordings in ChReef-injected Mongolian gerbils. a**, Exemplary oABR trace (representative of N=10 ChReef-injected animals). **b**, oABR thresholds computed across animals with a clear oABR responses, points represent individual gerbils (N=10). **c**, Characteristic frequencies (CF) as a function of recording depth. Dashed line: linear fit of all CFs for each animal, according to Pearson's correlation coefficient. Inset displays distribution of tonotopic slopes computed from linear fits across all animals. **d**, Example coronal brain slice of the inferior colliculus of a gerbil injected with ChReef-YFP (representative of N=2 brains) stained for Parvalbumin (PV, red) and GFP (green). **e**, Heatmap computed like in Fig. 5 for acoustic stimulation after kanamycin injection (no activity indicating deafening, representative of N=8 ChReef-injected and deafened animals).

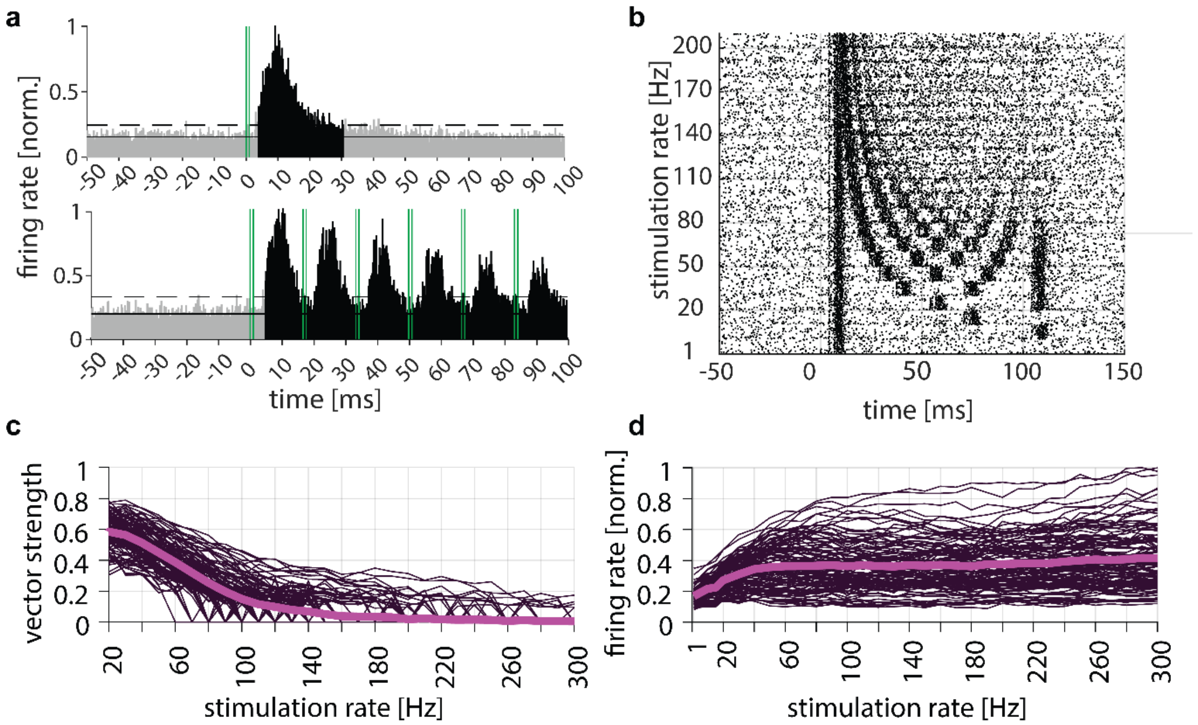


**Supplementary Figure 15. Temporal resolution of optogenetic activation of the auditory pathway by means of inferior colliculus recordings in Mongolian gerbils.**

a, Peri-Stimulus Time Histogram (PSTH) computing spike rates in bins of 0.2 ms recorded from stimulation at ~1 mW with an optical fiber emitting green light (522 nm) for single pulses (N = 10, top) and pulsing at 60Hz (N = 10 gerbils, bottom) normalized to the highest spike rate across animals. Green lines indicate stimulus timing. Dashed horizontal lines indicate 3x mean absolute deviation (MAD) used for thresholding and bins reaching threshold condition are highlighted in black. b, Exemplary spike raster computed from stimulating at 1 mW across different stimulation rates used for each of the 30 repetitions. Stimulus used were 100 ms pulse trains of 1 ms light pulses delivered at various stimulation rates. c-d, Individual traces computed across all animals and all responsive channels demonstrating vector strength (c) and firing rates (d) recorded and normalized to highest firing rate computed respectively. Average shown in bold magenta lines.

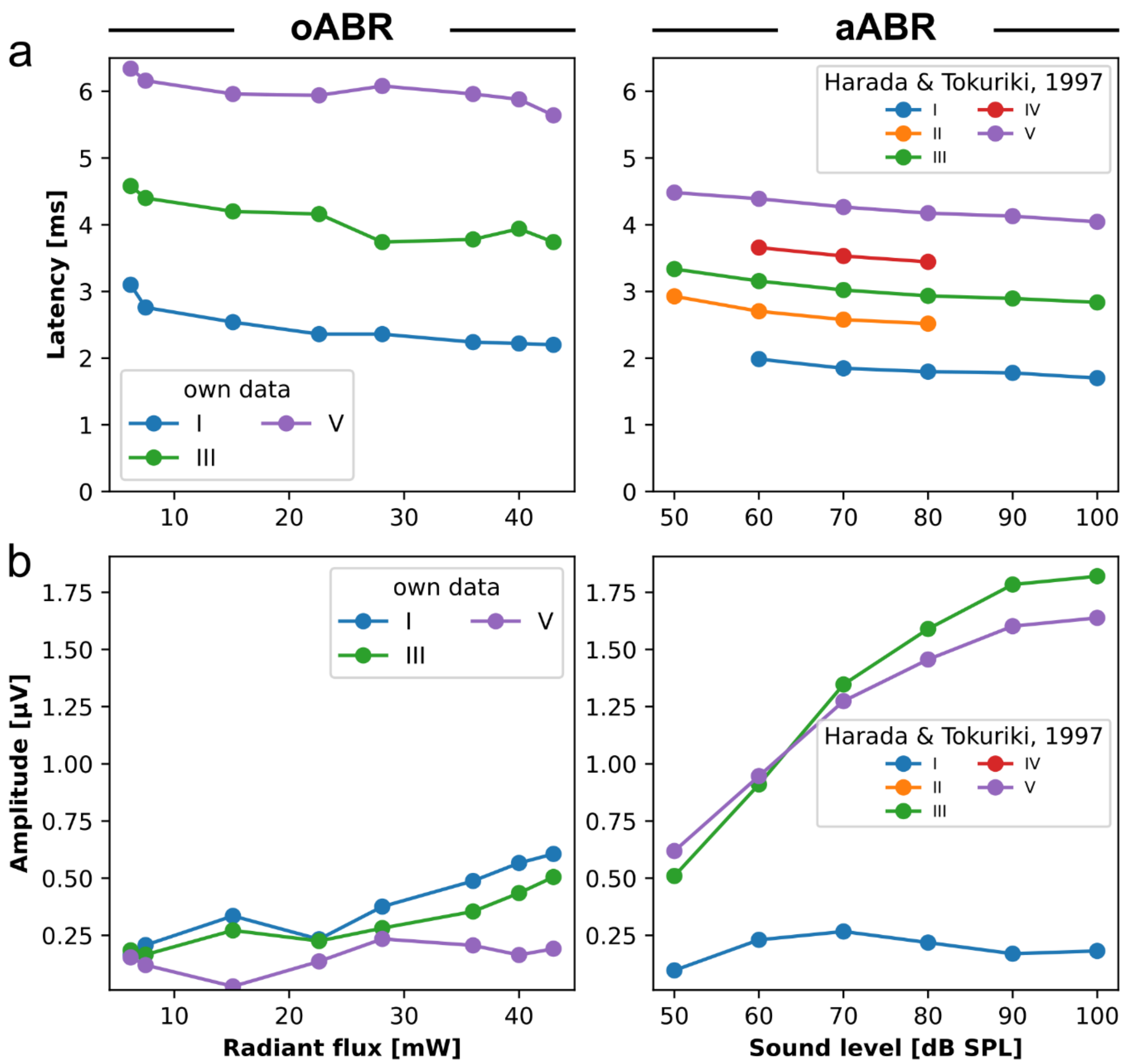


**Supplementary Figure 16. Comparison of oABR from Figure 5j and published aABR data.** oABR (left panels) and aABR (right panels) peaks were labelled according to their latency of occurence and their latency (**a**) as well as their amplitude (**b**) plotted as a function of stimulus intensity for the oABR positive animal (n = 1/9). Published aABR data (replotted from Harada & Tokuriki, 1997; their Fig. 3 and 5)[2] were gathered after stimulation with broadband clicks of varying sound level and presented at a similar repetition rate as oABR (10 Hz for aABR vs. 9 Hz for oABR).

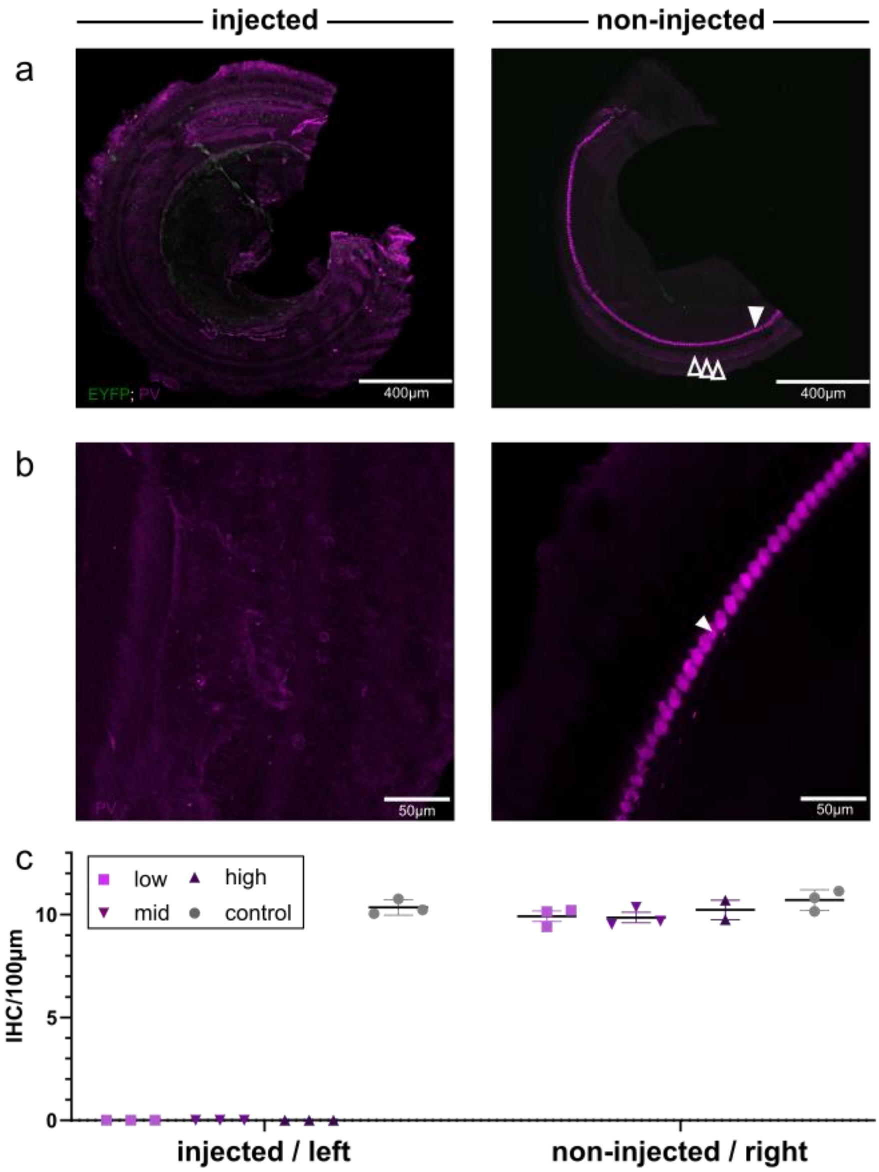


**Supplementary Figure 17. Quantification of inner hair cell loss following deafening of the injected left side in the common marmoset.**

**a,** Confocal images of cochlear whole mounts of an injected and fully deafened inner ear (left panel: 0 inner hair cells (IHC) observed) vs. the contralateral, non-injected side of the same animal (right panel: 9.678 cells/100µm). IHCs are indicated by a closed arrow head, outer hair cells by open arrow heads. **b,** Confocal images of zoomed-in ROI of the IHC (if present), same preparation as in a. **c,** IHC cell density across the across the different titer (n = 9) and control groups (n = 3). Note, that the IHC density on the non-injected, right side was comparable to IHC density in control animals (left: mean = 10.3 cells/100µm, right: mean = 10.7 cells/100µm).

**Supplementary Table 1. Stationary-Peak-Ratio, $EC_{50}$ values, stationary current densities and $\tau_{off}$ values of ChRmine variants.**

| ChR variant | Stationary-Peak-Ratio | $EC_{50}$ [µW/mm$^2$] | $J_{-60\ mV}$ [pA/pF] | $\tau_{off}$ [ms] |
|---|---|---|---|---|
| ChReef | 0.62 ± 0.15 | 20 ± 8 | 97.6 ± 65.0 | 58.3 ± 12.5 |
| ChRmine | 0.22 ± 0.12 | 14 ± 5 | 21.6 ± 15.8 | 63.5 ± 15.7 |
| ChRmine T218L | 0.44 ± 0.13 | 29 ± 15 | 64.8 ± 38.8 | 59.1 ± 21.3 |
| ChRmine S220A | 0.62 ± 0.14 | 31 ± 17 | 54.3 ± 27.2 | 152.7 ± 9.8 |

NG cells, transiently transfected with the shown ChRmine variants were investigated by whole-cell patch-clamp recordings at a membrane potential of -60 mV. Photocurrents were measured upon illumination with green light pulses (saturating intensity of 23 mW/mm$^2$, $\lambda$=532 nm) of 2 s for stationary current density determination and of 3 ms for $\tau_{off}$ determination. The stationary current densities ($J_{-60mV}$) were calculated as the quotient of the mean stationary current and the capacitance of the cell. The $\tau_{off}$ values were determined by a fit of the decaying photocurrent, immediately after cessation of the light pulse, to a monoexponential function. The half maximal activation ($EC_{50}$) was determined by hyperbolic fitting. For ChRmine wt the fitting range was restricted to the range, which could be approximated by a hyperbolic function. The stationary-peak-ratio was calculated as the quotient of the mean stationary current of the last 100 ms of the 2s light pulse and the peak current. Shown are mean and standard deviation (SD).

**Supplementary Table 2. Stationary current densities and $\tau_{off}$ values of ChR variants.**

| **ChR variant** | **$J_{-60\ mV}$ [pA/pF]** | **$\tau_{off}$ [ms]** |
|---|---|---|
| ChReef (a) | 97.6 ± 65.0 | 58.3 ± 12.5 |
| Catch (b) | 37.0 ± 12.5 | 34.4 ± 7.3 |
| ChR2 T159C (c) | 30.6 ± 18.4 | 28.3 ± 7.4 |
| Chronos (d) | 24.0 ± 7.5 | 3.5 ± 0.6 |
| Chrimson (e) | 24.0 ± 6.8 | 23.6 ± 1.5 |
| f-Chrimson (f) | 34.2 ± 12.7 | 5.7 ± 0.4 |

NG cells transfected with the shown ChR variants were investigated by whole-cell patch-clamp recordings at a membrane potential of -60 mV. Photocurrents were measured upon illumination with saturating light pulses of 0.5 s or 2 s for stationary current density determination and of 3 ms for $\tau_{off}$ determination. The stationary current densities ($J_{-60mV}$) were calculated as the quotient of the mean stationary current and the capacitance of the cell. The $\tau_{off}$ values were determined by a fit of the decaying photocurrent, immediately after cessation of the light pulse, to a monoexponential function. Shown are mean and standard deviation (SD).

**Supplementary Table 3. Exact (Mann-Whitney U-test) and adjusted (Bonferroni) p-values from the Figure 1a.**

| | protein1 | protein2 | p-value | adjusted p-value | significance |
|---|---|---|---|---|---|
| **0** | ChRmine | ChRmine S220A | 1,28E-14 | 7,65E-14 | **** |
| **1** | ChRmine | ChRmine T218L | 6,33E-07 | 3,8E-06 | **** |
| **2** | ChRmine | ChReef | 5,84E-09 | 3,51E-08 | **** |
| **3** | ChRmine S220A | ChRmine T218L | 7,09E-05 | 0,000426 | *** |
| **4** | ChRmine S220A | ChReef | 0,955443 | 1 | NS |
| **5** | ChRmine T218L | ChReef | 0,006471 | 0,038826 | * |

**Supplementary Table 4. Exact (Mann-Whitney U-test) and adjusted (Bonferroni) p-values from the Figure 1b.**

| | protein1 | protein2 | p-value | adjusted p-value | significance |
|---|---|---|---|---|---|
| 0 | Chronos | Chrimson | 0,365031116 | 1 | NS |
| 1 | Chronos | fChrimson | 0,010907843 | 0,392682347 | NS |
| 2 | Chronos | CatCh | 0,001362043 | 0,049033555 | * |
| 3 | Chronos | ChR2 T159C | 0,468800617 | 1 | NS |
| 4 | Chronos | ChRmine | 0,120278866 | 1 | NS |
| 5 | Chronos | ChReef | 6,47041E-06 | 0,000232935 | *** |
| 6 | Chronos | ChRmine S220A | 5,89845E-05 | 0,002123442 | ** |
| 7 | Chronos | ChRmine T218L | 0,000774258 | 0,027873292 | * |
| 8 | Chrimson | fChrimson | 0,012066365 | 0,43438914 | NS |
| 9 | Chrimson | CatCh | 0,012066365 | 0,43438914 | NS |
| 10 | Chrimson | ChR2 T159C | 0,480707528 | 1 | NS |
| 11 | Chrimson | ChRmine | 0,264624907 | 1 | NS |
| 12 | Chrimson | ChReef | 2,71935E-06 | 9,78965E-05 | **** |
| 13 | Chrimson | ChRmine S220A | 0,000147026 | 0,005292931 | ** |
| 14 | Chrimson | ChRmine T218L | 0,003350242 | 0,120608728 | NS |
| 15 | fChrimson | CatCh | 0,843831425 | 1 | NS |
| 16 | fChrimson | ChR2 T159C | 0,594853424 | 1 | NS |
| 17 | fChrimson | ChRmine | 0,00356568 | 0,128364494 | NS |
| 18 | fChrimson | ChReef | 0,000131121 | 0,004720342 | ** |
| 19 | fChrimson | ChRmine S220A | 0,005860854 | 0,210990738 | NS |
| 20 | fChrimson | ChRmine T218L | 0,012612069 | 0,454034489 | NS |
| 21 | CatCh | ChR2 T159C | 0,403317565 | 1 | NS |
| 22 | CatCh | ChRmine | 0,001150439 | 0,041415819 | * |
| 23 | CatCh | ChReef | 0,000286779 | 0,010324048 | * |
| 24 | CatCh | ChRmine S220A | 0,016623175 | 0,598434306 | NS |
| 25 | CatCh | ChRmine T218L | 0,023219344 | 0,835896385 | NS |
| 26 | ChR2 T159C | ChRmine | 0,13872372 | 1 | NS |
| 27 | ChR2 T159C | ChReef | 0,000614487 | 0,022121537 | * |
| 28 | ChR2 T159C | ChRmine S220A | 0,012341309 | 0,444287123 | NS |
| 29 | ChR2 T159C | ChRmine T218L | 0,016770456 | 0,603736421 | NS |
| 30 | ChRmine | ChReef | 4,38554E-08 | 1,5788E-06 | **** |
| 31 | ChRmine | ChRmine S220A | 1,90348E-08 | 6,85252E-07 | **** |
| 32 | ChRmine | ChRmine T218L | 3,95657E-06 | 0,000142436 | *** |
| 33 | ChReef | ChRmine S220A | 0,004073119 | 0,146632286 | NS |
| 34 | ChReef | ChRmine T218L | 0,108625377 | 1 | NS |
| 35 | ChRmine S220A | ChRmine T218L | 0,324126588 | 1 | NS |

**Supplementary Table 5. Exact (Student's t-test) and adjusted (Bonferroni) p-values from the Figure 1c.**

| | protein1 | protein2 | p-value | adjusted p-value | significance |
|---|---|---|---|---|---|
| **0** | Chronos | Chrimson | 1,63E-12 | 3,43286E-11 | **** |
| **1** | Chronos | fChrimson | 1,15E-05 | 0,000241192 | *** |
| **2** | Chronos | CatCh | 1,75E-09 | 3,66694E-08 | **** |
| **3** | Chronos | ChR2 T159C | 1,81E-07 | 3,8001E-06 | **** |
| **4** | Chronos | ChRmine | 7,15E-08 | 1,50064E-06 | **** |
| **5** | Chronos | ChReef | 1,28E-08 | 2,69656E-07 | **** |
| **6** | Chrimson | fChrimson | 7,1E-09 | 1,49005E-07 | **** |
| **7** | Chrimson | CatCh | 0,007169 | 0,150557442 | NS |
| **8** | Chrimson | ChR2 T159C | 0,189567 | 1 | NS |
| **9** | Chrimson | ChRmine | 0,000239 | 0,005013337 | ** |
| **10** | Chrimson | ChReef | 0,000113 | 0,002380895 | ** |
| **11** | fChrimson | CatCh | 8,63E-07 | 1,81211E-05 | **** |
| **12** | fChrimson | ChR2 T159C | 3,26E-05 | 0,000684998 | *** |
| **13** | fChrimson | ChRmine | 1,07E-05 | 0,000225602 | *** |
| **14** | fChrimson | ChReef | 3,07E-06 | 6,4388E-05 | **** |
| **15** | CatCh | ChR2 T159C | 0,20822 | 1 | NS |
| **16** | CatCh | ChRmine | 0,000198 | 0,004150877 | ** |
| **17** | CatCh | ChReef | 0,000183 | 0,003835063 | ** |
| **18** | ChR2 T159C | ChRmine | 7,83E-05 | 0,00164414 | ** |
| **19** | ChR2 T159C | ChReef | 6,58E-05 | 0,001380869 | ** |
| **20** | ChRmine | ChReef | 0,511891 | 1 | NS |

**Supplementary Table 6. Exact p-values (Tukey's HSD test) from the Figure 1i.**

| | protein1 | protein2 | p-value | significance |
|---|---|---|---|---|
| **0** | CatCh_stat | ChReef_ns | 2,51E-05 | **** |
| **1** | CatCh_stat | ChReef_stat | 0,00596 | ** |
| **2** | CatCh_stat | ChRmine_ns | 4,52E-07 | **** |
| **3** | CatCh_stat | ChRmine_stat | 0,001495 | ** |
| **4** | ChReef_ns | ChReef_stat | 0,61518 | NS |
| **5** | ChReef_ns | ChRmine_ns | 0,898008 | NS |
| **6** | ChReef_ns | ChRmine_stat | 0,904588 | NS |
| **7** | ChReef_stat | ChRmine_ns | 0,150454 | NS |
| **8** | ChReef_stat | ChRmine_stat | 0,987161 | NS |
| **9** | ChRmine_ns | ChRmine_stat | 0,41686 | NS |

**Supplementary Table 7. List of primers used for ChR mutant generation.**

| construct | template | sequence of forward primer | sequence of reverse primer |
|---|---|---|---|
| ChRmine F219Y | humanized ChRmine | 5’-GCCGTGTT TACCTACTCCA TGCTGTGG-3’ | 5’-CCACAGCA TGGAGTAGGTAA ACACGGC-3’ |
| ChRmine T218L | humanized ChRmine | 5´-GTTCGCCGTG TTTCTGTTCTCCA TGCTGTG-3´ | 5´-CACAGCATGG AGAACAGAAACAC GGCGAAC-3´ |
| ChRmine S220A | humanized ChRmine | 5´-GTGTTTACCTT CGCCATGCTG TGGATTC-3´ | 5´- GAATCCACAG CATGGCGAAG GTAAACAC -3´ |
| ChRmine T218L/S220A | humanized ChRmine T218L | 5´-CGTGTTTCTG TTCGCCATGCT GTGGATTCTG -3´ | 5´- CAGAATCCAC AGCATGGCGAA CAGAAACACG -3´ |
| ChR2 T159C | humanized ChR2 | 5´-TCCGACAT CGGCTGTATCGTAT GGGG-3´ | 5´- CCCCAT ACGATACAGCCG ATGTCGGA-3´ |
| ChRmine Y260F | humanized ChRmine | 5´-TGGCCA AGTCCTGTTTTG GCTTTGCCCTG-3´ | 5´-CAGGG CAAAGCCAAAACA GGACTTGGCCA-3´ |
| *Co*ChR L112C | humanized *Co*ChR | 5´-ACTTGTC CCGTCATCTGTATCC ACCTGAGCAAC-3´ | 5´-GTTGCTCA GGTGGATACAGA TGACGGGACAAGT-3´ |
| *Co*ChR H94E/L112C | humanized *Co*ChR L112C | 5´-TGGCTA ACGGAGAGCGAG TCCAGTGGC-3´ | 5´-GCCACTGG ACTCGCTCTCCGT TAGCCA-3´ |
| *Co*ChR H94E/L112C/ K264T | humanized *Co*ChR H94E/L112C | 5´- CGACATC CGGAAGACCACCA AGATCAACGTG-3´ | 5´-CACGTTG ATCTTGGTGGTC TTCCGGATGTCG-3´ |

**Supplementary Table 8. Solutions used for acute slice in vitro electrophysiology**

All compounds used are listed with their concentrations. QX314 stands for N-(2,6-dimethylphenylcarbamoylmethyl) triethylamonium chloride, an intracellular sodium channel blocker.

| *aCSF 2mM $Ca^{2+}$* | | *Intracellular solution* | |
|---|---|---|---|
| **Compound Name (Source)** | **Concentration (mM)** | **Compound name (Source)** | **Concentration (mM)** |
| Glucose (Merck) | 13 | K-gluconate (Merck) | 108 |
| NaCl (Merck) | 125 | HEPES (Merck) | 9 |
| $CaCl_2$ (Merck) | 2 | EGTA (Merck) | 9 |
| $MgCl_2$ (Merck) | 1 | $Na_2$Phosphocreatine (Merck) | 14 |
| $NaHCO_3$ (Merck) | 26 | ATP-Mg (Merck) | 4 |
| $NaH_2PO_4$ * $H_2O$ (Merck) | 1.25 | GTP-Na (Merck) | 0.3 |
| KCl (Merck) | 2.5 | $MgCl_2$ (Merck) | 4.5 |
| Na-Pyruvate (Merck) | 2 | QX-314 (Merck) | 1 |
| Na L-ascorbate (Merck) | 0.7 | | |

*Cutting Solution*

| **Compound Name (Source)** | **Concentration (mM)** |
|---|---|
| Glucose (Merck) | 20 |
| Sucrose (Merck) | 120 |
| NaCl (Merck) | 50 |
| $CaCl_2$ (Merck) | 0.2 |
| $MgCl_2$ (Merck) | 6 |
| $NaHCO_3$ (Merck) | 26 |
| $NaH_2PO_4$ * $H_2O$ (Merck) | 1.25 |
| KCl (Merck) | 2.5 |
| Na-Pyruvate (Merck) | 2 |
| Na L-ascorbate (Merck) | 0.7 |
| Na L-Lactate (Merck) | 3 |


1. Dieter, A., Duque-Afonso, C. J., Rankovic, V., Jeschke, M. & Moser, T. Near physiological spectral selectivity of cochlear optogenetics. *Nature Communications* **10**, 1962 (2019).
2. Harada, T. & Tokuriki, M. Effects of click intensity and frequency on the brain-stem auditory evoked potentials in the common marmoset (Callithrix jacchus). *Journal of Veterinary Medical Science* **59**, 561–567 (1997).

**Supplementary Table 9. Exact (Mann-Whitney U-test) and adjusted (Bonferroni) p-values from the Extended Data Figure 2b.**

| | protein1 | protein2 | p-value | adjusted p-value | significance |
|---|---|---|---|---|---|
| **0** | ChRmine | ChReef | 1,22E-08 | 1,22E-07 | **** |
| **1** | ChRmine | ChRmine T218L | 8,66E-05 | 0,000866 | *** |
| **2** | ChRmine | ChRmine S220A | 0,040016 | 0,400158 | NS |
| **3** | ChRmine | ChRmine F219Y | 6,52E-11 | 6,52E-10 | **** |
| **4** | ChReef | ChRmine T218L | 0,038726 | 0,38726 | NS |
| **5** | ChReef | ChRmine S220A | 0,00321 | 0,032098 | * |
| **6** | ChReef | ChRmine F219Y | 2,72E-11 | 2,72E-10 | **** |
| **7** | ChRmine T218L | ChRmine S220A | 0,271432 | 1 | NS |
| **8** | ChRmine T218L | ChRmine F219Y | 5,36E-10 | 5,36E-09 | **** |
| **9** | ChRmine S220A | ChRmine F219Y | 4,96E-09 | 4,96E-08 | **** |

**Supplementary Table 10. Exact (Mann-Whitney U-test) and adjusted (Bonferroni) p-values from the Extended Data Figure 2c.**

| | protein1 | protein2 | p-value | adjusted p-value | significance |
|---|---|---|---|---|---|
| **0** | GtCCR4 | ChRmine | 3,34E-10 | 9,35E-09 | **** |
| **1** | GtCCR4 | GtCCR2 | 0,338786 | 1 | NS |
| **2** | GtCCR4 | CoChR-3M | 1,31E-09 | 3,66E-08 | **** |
| **3** | GtCCR4 | CatCh | 1,44E-07 | 4,02E-06 | **** |
| **4** | GtCCR4 | ChReef | 1,57E-10 | 4,4E-09 | **** |
| **5** | GtCCR4 | ChRmine Y260F | 1,52E-09 | 4,26E-08 | **** |
| **6** | GtCCR4 | CoChR | 1,59E-10 | 4,46E-09 | **** |
| **7** | ChRmine | GtCCR2 | 2,16E-13 | 6,05E-12 | **** |
| **8** | ChRmine | CoChR-3M | 5,88E-06 | 0,000165 | *** |
| **9** | ChRmine | CatCh | 1,58E-06 | 4,43E-05 | **** |
| **10** | ChRmine | ChReef | 1,22E-08 | 3,41E-07 | **** |
| **11** | ChRmine | ChRmine Y260F | 0,743772 | 1 | NS |
| **12** | ChRmine | CoChR | 0,071397 | 1 | NS |
| **13** | GtCCR2 | CoChR-3M | 2,41E-12 | 6,76E-11 | **** |
| **14** | GtCCR2 | CatCh | 4,15E-11 | 1,16E-09 | **** |
| **15** | GtCCR2 | ChReef | 5,45E-14 | 1,52E-12 | **** |
| **16** | GtCCR2 | ChRmine Y260F | 3,12E-12 | 8,75E-11 | **** |
| **17** | GtCCR2 | CoChR | 7,75E-15 | 2,17E-13 | **** |
| **18** | CoChR-3M | CatCh | 1,08E-10 | 3,03E-09 | **** |
| **19** | CoChR-3M | ChReef | 0,363146 | 1 | NS |
| **20** | CoChR-3M | ChRmine Y260F | 2,49E-06 | 6,97E-05 | **** |
| **21** | CoChR-3M | CoChR | 1,22E-08 | 3,42E-07 | **** |
| **22** | CatCh | ChReef | 2,91E-13 | 8,15E-12 | **** |
| **23** | CatCh | ChRmine Y260F | 5,09E-06 | 0,000143 | *** |
| **24** | CatCh | CoChR | 0,000283 | 0,007921 | ** |
| **25** | ChReef | ChRmine Y260F | 4,51E-09 | 1,26E-07 | **** |
| **26** | ChReef | CoChR | 1,03E-12 | 2,9E-11 | **** |
| **27** | ChRmine Y260F | CoChR | 0,175989 | 1 | NS |

**Supplementary Table 11. Exact p-values (Tukey's HSD test) from the Extended Data Figure 2d.**

| | protein1 | protein2 | p-value | significance |
|---|---|---|---|---|
| **0** | CatCh | ChReef | 0,346598051 | NS |
| **1** | CatCh | ChRmine | 0,000225641 | *** |
| **2** | CatCh | ChRmine Y260F | <1E-13 | **** |
| **3** | CatCh | CoChR | 0,998691665 | NS |
| **4** | CatCh | CoChR-3M | <1E-13 | **** |
| **5** | CatCh | GtCCR2 | 0,022199136 | * |
| **6** | CatCh | GtCCR4 | 0,703629469 | NS |
| **7** | ChReef | ChRmine | 0,27132902 | NS |
| **8** | ChReef | ChRmine Y260F | 7,63148E-09 | **** |
| **9** | ChReef | CoChR | 0,159017706 | NS |
| **10** | ChReef | CoChR-3M | <1E-13 | **** |
| **11** | ChReef | GtCCR2 | 2,73628E-06 | **** |
| **12** | ChReef | GtCCR4 | 0,018664277 | * |
| **13** | ChRmine | ChRmine Y260F | 0,000402241 | *** |
| **14** | ChRmine | CoChR | 0,000127347 | *** |
| **15** | ChRmine | CoChR-3M | <1E-13 | **** |
| **16** | ChRmine | GtCCR2 | 6,73428E-12 | **** |
| **17** | ChRmine | GtCCR4 | 1,55577E-05 | **** |
| **18** | ChRmine Y260F | CoChR | 2,2593E-13 | **** |
| **19** | ChRmine Y260F | CoChR-3M | <1E-13 | **** |
| **20** | ChRmine Y260F | GtCCR2 | <1E-13 | **** |
| **21** | ChRmine Y260F | GtCCR4 | 4,16334E-13 | **** |
| **22** | CoChR | CoChR-3M | <1E-13 | **** |
| **23** | CoChR | GtCCR2 | 0,22726708 | NS |
| **24** | CoChR | GtCCR4 | 0,960896094 | NS |
| **25** | CoChR-3M | GtCCR2 | <1E-13 | **** |
| **26** | CoChR-3M | GtCCR4 | <1E-13 | **** |
| **27** | GtCCR2 | GtCCR4 | 0,974017951 | NS |

#### Supplementary Notes

##### Supplementary Note 1: Cell culture and transfection

The manual patch-clamp recordings of ChR variants were conducted in the neuroma glioblastoma cell line NG108-15 (ATCC, HB-12377TM, Manassas, USA) cultured in Dulbecco's Modified Eagle Medium (DMEM, Sigma, St. Louis, USA) supplemented with 10 % fetal calf serum (Sigma, St. Louis, USA) and 1 % penicillin/streptomycin (Sigma, St. Louis, USA) (supplemented DMEM: DMEM$^+$) at 37 °C and 5 % $CO_2$. Cells were seeded on 24-well plates one day prior to transfection by Lipofectamine with pcDNA3.1(-) derivatives carrying the aforementioned ChR variants at a NG108-15 cell confluency of 50-70 %. For each well, a transfection mix of 100 µl DMEM, 3 µl Lipofectamine LTX (Invitrogen, Carlsbad, USA) and 500 ng of the pcDNA3.1(-) derivatives was prepared and added to a well with 400 µl of DMEM$^+$. Twenty-four hours after transfection the medium was exchanged against 500 µl DMEM$^+$ supplemented with 1 µM all-trans retinal.

For automated patch-clamp recordings and automated spinning-disc confocal microscopy, HEK293T cells (DSMZ, Braunschweig, Germany) or NG108-15 (ATCC, HB-12377TM, Manassas, USA) cells were seeded in a T75 culture flasks or 6-well plates and allowed to grow for one day before transfection. The cells were seeded at a density of 16 thousands cells/cm$^2$ in DMEM$^+$ (initial concentration before adhesion 80 thousands cells/ml). For noise analysis experiments transfection of HEK293T cells was done in T75 flasks. Immediately prior to transfection, the media was replaced with 12 mL of OptiMEM (ThermoFisher, Waltham, Massachusetts, U.S.) with 1.25 µM all-trans-retinal (Sigma, St. Louis, MO, U.S.). Then, 19.5 µg of plasmid DNA and 78.1 µg of polyethyleneimine (PEI, Polysciences, Warrington, PA, U.S.) were separately dissolved in 1.5 ml of OptiMEM (ThermoFisher, Waltham, Massachusetts, U.S.) each. The DNA and PEI solutions were then combined and incubated for 15 minutes at room temperature. The resulting DNA/PEI mix was added to the cells and incubated at 37°C and 5% $CO_2$ for 4 hours. After transfection, the media in the flask was changed back to $DMEM^+$ supplemented with 1 μM all-trans-retinal. The cells were then incubated at 37°C and 5% $CO_2$ for 40 hours prior to the automated patch-clamp experiment. For all other experiments with automated patch-clamp transfections of HEK293T or NG108-15 cells were done in 6-well plates without prior media exchange. For each well 2.5 μg of plasmid DNA and 7.5 μg of PEI were separately dissolved in 0.2 ml of OptiMEM. The DNA and PEI solutions were then combined and incubated for 15 minutes at room temperature. The resulting DNA/PEI mix was added to the wells of 6-well plate and incubated at 37°C and 5% $CO_2$ for 16 hours. After transfection, the media in the flask was exchanged to new $DMEM^+$ supplemented with 1 μM all-trans-retinal. The cells were then incubated at 37°C and 5% $CO_2$ for 28 hours prior to the automated patch-clamp experiment. For automated spinning-disc confocal microscopy NG108-15 cell transfections were done in 96-well glass bottom microscopy grade plates (Greiner, Sensoplate microplate 96 well, 655892). For each well 83 ng of plasmid DNA and 0.25 μg of PEI were separately dissolved in 7 μl of OptiMEM. The DNA and PEI solutions were then combined and incubated for 15 minutes at room temperature. The resulting DNA/PEI mix was added to the wells of the 96-well plate and incubated at 37°C and 5% $CO_2$ for 16 hours. After transfection, the media in the flask was exchanged to new $DMEM^+$ supplemented with 1 μM all-trans-retinal. The cells were then incubated at 37°C and 5% $CO_2$ for 28 hours prior to the automated spinning-disc confocal microscopy experiment.

**Supplementary Note 2: Protocols for manual patch-clamp recordings and data analysis**

Patch pipettes with a resistance of 2-6 MΩ were fabricated from thin-walled borosilicate glass on a horizontal puller (Model P-1000, Sutter Instruments, Novato, USA). The series resistance was <15 MΩ. Light pulses were applied by a fast computer-controlled shutter (Uniblitz LS6ZM2, Vincent Associates, Rochester, USA) using diode-pumped solid-state lasers focused into an optic fiber (d=400 μm). The open-source statistic software "R" as well as Origin 9.0

(OriginLab, Inc., Northampton, MA, USA) and GraphPad Prism (GraphPad Software, La Jolla, CA, USA) were employed for data analysis.

The photocurrents of the ChRmine variants were measured in response to green light pulses ($\lambda$=532 nm) of varying duration with a saturating intensity of 23 mW/mm$^2$. The photocurrents of Chronos, Catch and ChR2 T159C were measured upon the application of blue light pulses ($\lambda$=473 nm) of varying duration with a saturating intensity of 31 mW/mm$^2$. The stationary current densities ($J_{-60\ mV}$) of the ChRmine variants were calculated as the quotient of the mean stationary current upon stimulation with a 2 s light pulse and the capacitance of the cell. The stationary current densities ($J_{-60\ mV}$) of Chronos, Catch and ChR2 T159C were calculated as the quotient of the mean stationary current upon stimulation with a 0.5 s light pulse and the capacitance of the cell. To avoid an experimental bias, cells were chosen for the recordings independent of the brightness of their fluorescent signal.

The stationary-peak-ratio of the photocurrents of the ChRmine variants was calculated as the quotient of the mean stationary current upon stimulation with a 2 s light pulse and the peak current. The peak recovery of the ChRmine variants was assessed by photocurrent recordings upon illumination with two subsequent 3 s light pulses with a varying time between the light pulses ($\Delta$t between pulses, ms, ranging from 50 to 3350 ms). Peak recovery was calculated as the quotient of the difference between the peak and stationary current of pulse two ($I_{p2}$-$I_{s2}$) and peak and stationary current of pulse one ($I_{p1}$-$I_{s1}$) in %. The time constant of the peak recovery ($\tau_{peak\text{-}recovery}$) was determined by a fit of the quotient of the difference between the peak and stationary currents of two subsequent 3s light pulses at saturating intensity to a monoexponential function.

The $\tau_{off}$ values of all shown ChR variants were determined by a fit of the decaying photocurrent elicited in response to a 3 ms light pulse to a monoexponential function. In order to investigate the dependence of the off-kinetics on the membrane potential $\tau_{off}$ values were determined at membrane potentials ranging from -120 mV to +60 mV.

The action spectra of the ChRmine variants were determined by peak current recordings in response to ns-light-pulses of wavelengths ranging from 430 to 590 nm with pulse energies at the different wavelengths set to equal photon counts of $5*10^{18}$ photons/$m^2$ using the Opolette 355 tunable laser system (Opotek Inc, Carlsbad, USA).

**Supplementary Note 3: Automated patch clamping procedure**

To begin, the S-type chip (Nanion, Munich, Germany) was filled with 80 µl per well of Divalent-Free solution, and the chamber's bottom was subsequently filled with Internal solution. The Divalent Free solution contained 140 mM NaCl, 4 mM KCl, and 10 mM HEPES, with pH adjusted to 7.4 using NaOH. The osmolarity of the solution was set to 289 mOsm by adjusting the concentration of glucose. The Internal solution was composed of 10 mM NaCl, 10 mM KCl, 110 mM KF, 10 mM HEPES, and 10 mM EGTA, with pH adjusted to 7.2 using KOH. The osmolarity of the solution was set to 285 mOsm by adjusting the concentration of glucose. Next, 20 µl per well of solution was removed, and 20 µl per well of detached cells were introduced to the chip. The cells were then captured by a suction pulse of -200 mbar for 5 s, followed by consecutive suction pulses of -50 mbar for 1 min. Subsequently, the voltage was decreased incrementally from 0 mV to -100 mV in steps of 25 mV while maintaining the cells under a pressure of -50 mbar HEK293T) or -80 mbar (NG108-15). After this, 20 µl per well of solution was removed from the chip, and 40 µl per well of seal enhancer solution was added. The seal enhancer solution contained 130 mM NaCl, 4 mM KCl, 10 mM $CaCl_2$, 1 mM $MgCl_2$, and 10 mM HEPES, with pH adjusted to 7.4 using NaOH. The osmolarity of the solution was set to 302 mOsm by adjusting the concentration of glucose. The chip then underwent three wash steps, during which 40 µl of solution was removed from each well and replaced with 40 µl per well of standard external solution (unless another extracellular solution composition is specified). After the second wash step, two suction pulses of -220 mbar were applied for 2 s each to establish a whole-cell configuration for HEK293T cells or three suction pulses of -350 mbar were applied for 5 s each to establish a whole-cell configuration for NG108-15 cells. After completing all of the procedures described, the remaining volume of solution in the wells was 100 μl.

**Supplementary Note 4: Cell treatment prior automated patch-clamp experiments**

The cells that were transfected were washed twice with DPBS (ThermoFisher, Waltham, Massachusetts, U.S.) and then treated with 3 ml of TrypLE (ThermoFisher, Waltham, Massachusetts, U.S.) at 37°C for 7 minutes (HEK293T) or 15 minutes (NG108-15). After the TrypLE treatment, 3 ml of standard external solution was added to the cells and incubated at 4°C for 5 minutes. The standard external solution was composed of 140 mM NaCl, 4 mM KCl, 2 mM $CaCl_2$, 1 mM $MgCl_2$, and 10 mM HEPES, with pH adjusted to 7.4 using NaOH. The osmolarity of the solution was set to 298 mOsm by adjusting the concentration of glucose. The cells were then resuspended by pipetting and centrifuged at 200 g for 10 minutes, and the supernatant was discarded. Finally, the cells were resuspended in 10 mL of Standard External Solution.

**Supplementary Note 5: Illumination in automated patch clamp experiment**

To illuminate the cells during the planar patch clamp experiment, a specialized illumination unit was used. The unit consists of 96 LEDs (Lumileds Luxeon Z, Schiphol, Haarlemmermeer, The Netherlands) that are coupled to light fibers. During the experiment, the end of the fiber is submerged under the solution and positioned 7 mm from the cell being measured. The illumination units with LEDs of different colors were used in the planar patch-clamp experiments: green LEDs ($\lambda_{max}$ = 530 nm, LXZ1-PM01), blue LEDs ($\lambda_{max}$ = 470 nm, LXZ1-

PB01), orange LEDs ($\lambda_{max}$ = 590 nm, LXZ1-PL02) and red LEDs ($\lambda_{max}$ = 632 nm, LXZ1-PD01). To regulate the intensity of light used in each measurement, the forward current of the illumination unit was adjusted. The intensity was measured directly at the output of the light fibers using a bolometer (Coherent OP-2 VIS, Santa Clara, California, U.S.). The intensity of light was specified for each type of measurement.

**Supplementary Note 6: Noise analysis**

The data collection for the noise analyses of both types started with measurement of photocurrent dependence on the membrane voltage (IV curve). The range of voltages was from -120 mV to 60 mV with equidistant 20 mV steps. Each sweep was separated into three consecutive steps, with 2 s in the dark, 2 s under the illumination of a saturating intensity for the stationary photocurrents of the measured ChRs (4 mW/mm$^2$ for ChReef and ChRmine; 6 mW/mm$^2$ for CatCh) and 1 s in the dark. This data was used to calculate the reversal potentials, which were used to determine the electrochemical potentials on the membranes. This step was followed by different protocols and data treatment procedures for stationary and non-stationary analyses. The data treatment encompassed a thorough selection of recordings that met our strict quality control criteria, followed by the processing of the selected raw traces using our proprietary Python library and custom scripts developed in-house.

For stationary analyses, all cells were held at a voltage of -100 mV or -60 mV. 40 consecutive pairs of sweeps, 6.5 s long each, were recorded at a sampling rate of 20 kHz. Each pair of sweeps was followed by 25 s of non-recorded waiting time in the dark. The first sweep of each pair had a 6-second illumination pulse at an intensity of 0.4 mW/mm$^2$ (0.6 mW/mm$^2$ for CatCh), and the second sweep was recorded under dark conditions.

Sweeps suitable for stationary noise analysis were selected based on the following criteria. Basic parameters of the seals were assessed for each sweep, and only sweeps with a seal resistance over 1 GΩ, a membrane capacitance lower than 50 pF, and a series resistance below 20 MΩ were used for further analysis. The parameters of the dark noise were then assessed using parts recorded under the dark conditions for the sweeps with data recorded under illumination conditions. Sweeps with a root mean square deviation (IRMS) of dark currents over 5.5 pA were disregarded. The stability of the noise was assessed by dividing the dark part of the current into 100 pieces, calculating IRMS in all of them, and disregarding sweeps with a standard deviation of IRMS over 0.3 pA. The recorded sweeps were grouped back into pairs of consecutive traces that were recorded under light and dark conditions. Only those pairs with stationary photocurrents over 200 pA during the interval from 2 to 6 s after turning on the illumination were considered. Cells that had four or more pairs of traces remaining after applying all of the aforementioned quality control criteria were selected for stationary noise analysis. For each pair, the interval from 2 to 6 s was taken. The sweep recorded under the light conditions was fitted to a double exponential function, and the resulting fit was subtracted from the sweep. A mean current was subtracted from the sweep recorded in the dark. After these corrections, the power spectral density (PSD) was calculated for both sweeps. The differences between the PSD under the light and dark conditions were determined and then averaged for each cell. The averaged difference PSDs were then fitted to the single Lorentzian function in a range from 2.5 to 25 Hz. This allowed for the determination of the corner frequency and the difference PSD at 0 Hz ($f_c$ and $S_0$ correspondingly as in eq. 2). Open probabilities of the channels ($P_o$) were then determined for each cell, taking into consideration their relation (eq. 3) to corner frequency and off kinetics (where $\tau_{off}$ was determined from the single exponential fit of the photocurrent decrease after turning off the light). Only cells whose difference PSD met the criteria of 1 Hz < $f_c$ < 15 Hz and 0 < $P_o$ < 1 were considered in the determination of the single channel conductance values.

$$S(f) = \frac{S_0}{1 + \left(\frac{f}{f_c}\right)^2} \qquad \text{[Equation 2]}$$

$$1 - P_o = \frac{1}{2\,\pi f_c \tau_{off}} \qquad \text{[Equation 3]}$$

The variances $\sigma^2$ of the noise associated with the opening and closing of the ChRs were determined by using the obtained values of $f_c$ and $S_0$ (eq. 4). The unitary photocurrents were subsequently calculated by applying the relation between the mean and variance in the binomial distribution (eq. 5) to the mean net stationary current I and $\sigma^2$. Afterwards, the unitary photocurrents were converted to single channel conductances by dividing them by the electrochemical potential gradient.

$$\sigma^2 = \frac{\pi\, f_c\, S_0}{2} \qquad \text{[Equation 4]}$$

$$i = \frac{\sigma^2}{I\,(1 - P_o)} \qquad \text{[Equation 5]}$$

In the non-stationary analysis, 300 sweeps were recorded for each cell at the holding voltage of -100 mV or -60 mV. At -100 mV each sweep had a 5 ms illumination step at an intensity of 14 mW/mm$^2$ forward current so that recordings are done close to single turnover conditions, preventing protein from going to the putative second open state upon absorption of a second photon. At -60 mV each sweep had an illumination step of 150 ms at an intensity of 0.4 mW/mm$^2$. After each illumination step the dark currents were recorded for 5 s.

The non-stationary noise analysis involved several steps to select the suitable sweeps for analysis. Firstly, basic parameters of the seals were assessed, and only those sweeps that met the following criteria were used: seal resistance over 1 GΩ, membrane capacitance lower than 50 pF, and series resistance below 20 MΩ. Sweeps with root mean square deviation (IRMS) of dark currents over 4.5 pA were disregarded. Only cells with a peak photocurrent over 500 pA (over 100 pA for -60 mV traces) were considered. To minimize possible unwanted effects of the photocurrent rundown, only the sets of at least 50 consecutive recordings (25 recordings for -60 mV traces) with negligible rundown of the peak photocurrent (lower than 1 pA/sweep) were considered and the variance of the photocurrent was determined as a half of the average of the squared differences between pairs of consecutive recordings. The dependence of the variance on the photocurrent value was then fitted linearly to determine the unitary photocurrent as a proportionality factor (eq. 6) and the background current variance as the free term ($\sigma_b^2$, squared IRMS at photocurrent of 0 pA). Finally, any cells that had linear fits indicating $\sigma_b$ smaller than 4 pA were excluded from the analysis, since none of the recordings showed such a low IRMS value based on its direct calculation in the dark current. The remaining cells were used to compute the single channel conductance.

$$\sigma^2 = P_o(1 - P_o)Ni^2 + \sigma_b^2 \approx P_oNi^2 + \sigma_b^2 = I \cdot i + \sigma_b^2 \qquad \text{[Equation 6]}$$

**Supplementary Note 7: Comparison of basic biophysical properties of ChRs measured with automated patch clamp: photocurrent densities, stationary to peak ratios and off kinetics**

The photocurrents of the ChR variants at a membrane voltage of -60 mV were measured in response to green LED pulses (λ=530 nm) for ChRmine variants and GtCCR4, or blue LED pulses (λ=470 nm) for CatCh, CoChR, CoChR-3M, and GtCCR2. Light pulses of 2 s duration with a saturating intensity (green LED at 5.6 mW/mm² and blue LED at 8.3 mW/mm²) were used to determine photocurrent densities and desensitization kinetics. The stationary photocurrent densities of the ChR variants were calculated as the quotient of the mean stationary photocurrent upon stimulation with a 2 s light pulse and the capacitance of the cell. Due to the lack of a fluorescence control, we selected only cells with peak photocurrents greater than 20 pA. To avoid bias that could inflate the apparent mean value of photocurrent density for small photocurrent variants, cells with photocurrent densities in the lowest 15% were excluded for each variant. Desensitization kinetics were determined by fitting the decays of the photocurrents from their peak to stationary levels to a biexponential function. Normalized values for fast and slow components' amplitudes were calculated as quotients of the corresponding amplitudes and the sum of the two amplitudes.

Light pulses of 5 ms duration with an intensity of 13.0 mW/mm² for the green LED (ChRmine variants, GtCCR4) and 19.3 mW/mm² for the blue LED (CatCh, CoChR, CoChR-3M, and GtCCR2) were used to determine off-kinetics. The $\tau_{off}$ values of all ChR variants were determined by fitting the decaying photocurrent elicited in response to a 5 ms light pulse to a monoexponential function. To determine $\tau_{off}$ for CoChR, only cells with stationary photocurrent densities lower than 50 pA/pF were selected to avoid effects of intracellular proton accumulation affecting pH-dependent off-kinetics. To investigate the dependence of the off-kinetics on the membrane potential, $\tau_{off}$ values were determined at membrane potentials ranging from -120 mV to +60 mV.

#### Supplementary Note 8: Intensity dependence of photocurrents measured with automated patch clamp

For intensity dependence experiments, light pulses of 2 s duration were used for green light (λ=530 nm) and orange light (λ=590 nm), while light pulses of 4 s duration were used for red light (λ=632 nm). The intensity of light was regulated by adjusting the forward current of the respective illumination unit. Alternating between fixed and variable light intensities was employed to control for photocurrent rundown effects. The stationary photocurrents recorded at varying intensities were normalized against preceding and subsequent measurements at fixed intensities. The resulting dependencies for each recorded cell were fitted to hyperbolic functions and normalized by their respective amplitudes from the fits. The normalized dependencies were then averaged between the cells.

**Supplementary Note 9: Relative ion permeability determination**

For the determination of relative ion permeability, photocurrents of ChRmine and ChReef in response to green LED pulses (λ=530 nm) of 2 s duration were measured at different voltages, and reversal potentials were determined. Permeability ratios were calculated according to the Goldman–Hodgkin–Katz equation. To assess the relative permeability of potassium ions relative to sodium (PK/PNa), we measured photocurrent at voltages ranging from -120 mV to +60 mV in 10 mV steps and assessed reversal potential shifts after exchanging the standard intracellular solution (KF-based) with a NaF-based intracellular solution containing 10 mM NaCl, 10 mM KCl, 110 mM NaF, 10 mM HEPES, and 10 mM EGTA, with pH adjusted to 7.2 using NaOH. A reverse exchange of the intracellular solution was performed to account for reversal potential drift. The shift in reversal potential due to the solution exchange was calculated as the difference between the reversal potential in the NaF-based intracellular solution and the average of the two reversal potentials measured in the KF-based intracellular solution (before exchange and after reverse exchange). Cells with a reversal potential drift greater than 5 mV were excluded from the analysis.

To assess the relative permeability of calcium ions relative to sodium (PCa/PNa), we measured photocurrent at voltages ranging from -120 mV to +60 mV in 10 mV steps and assessed reversal potential shifts after extracellular solution dilution, which changed the NaCl concentration from [NaCl]_ini = 140 mM to [NaCl]_fin = 11 mM and the $CaCl_2$ concentration from [$CaCl_2$]_ini = 2.5 mM to [$CaCl_2$]_fin = 83 mM. An additional series of extracellular solution dilutions was performed to return sodium and calcium concentrations to their initial values to account for reversal potential drift. The shift in reversal potential due to the solution exchange was calculated as the difference between the reversal potential in the Ca-based extracellular solution and the average of the two reversal potentials measured in the Na-based extracellular solution (before dilution and after reverse dilution series). Cells with a reversal potential drift greater than 10 mV were excluded from the analysis.

To assess the relative permeability of protons relative to sodium ($P_H/P_{Na}$), we measured photocurrent at voltages ranging from -121 mV to 59 mV in 3 mV steps. The permeability of protons relative to sodium was calculated from the photocurrent reversal potential using a standard intracellular solution and an NMG-based extracellular solution containing 90 mM NMDG, 3.2 mM NaCl, and 10 mM MES, with pH adjusted to 6.4 using HCl.

**Supplementary Note 10: Staining and data acquisition for automated spinning disc confocal microscopy**

To assess the subcellular localization of ChRs in NG108-15 cells, the transfected cells were stained with Hoechst 33342 (1:2000, Invitrogen, H3570) and CellTracker Deep Red (1:1000, Invitrogen, 2521042). The culture medium was aspirated, and the cells were washed with 50 µl of DPBS. Afterward, 50 µl of Opti-MEM containing the dyes was added to the cells, followed by incubation at 37 °C for 30 minutes. Post incubation, the cells were washed again with DPBS and 200 µl of DMEM$^{+}$ was added. The stained cells were then imaged using the CellVoyager CQ1 microscope (Yokogawa). Identical imaging parameters were employed across all experiments. A 40x/0.95 dry objective (UPLXAPO40X) was used for acquisitions. Hoechst 33342 fluorescence was excited with a 405 nm laser at 20% power and a 500 ms exposure time, using a BP447/60 emission filter. EYFP fluorescence was excited with a 488 nm laser at 20% power and a 500 ms exposure time, using a BP525/50 emission filter. CellTracker Deep Red was excited with a 640 nm laser at 20% power and a 500 ms exposure time, using a BP685/40 emission filter. Thereby an autofocus routine in the Hoechst channel ensured imaging close to the central planes of the cells.

**Supplementary Note 11: Analysis of spinning disc microscopy data**

We wrote a Python script for fluorescence line profile analysis. First, the program identified the centroids of the nuclei in the Hoechst channel using the mean-shift clustering algorithm with a bandwidth of 8.125 µm. These centroids were then used to generate 2D square crops of the images, each containing one complete NG cell. Cell membrane masks were automatically generated based on the cytoplasmic fluorescence from the CellTracker channel, using a threshold derived from a previously measured calibration curve. This curve represented the relationship between CellTracker fluorescence intensity at the cell border and the average foreground fluorescence. For calibration, ChRmine-TS-EYFP-ES expressing cells which showed predominant plasma membrane-localization were manually selected. The foreground fluorescence in the CellTracker channel was averaged in regions where the CellTracker fluorescence exceeded that of the nucleus, ensuring it remained above background noise. The optimal CellTracker fluorescence threshold was determined by maximizing the average EYFP fluorescence along the cell borders, calculated using various CellTracker thresholds. Importantly, the dataset that was used for calibration curve generation was not included into the line profile analysis presented in the paper. The line profiles were automatically positioned perpendicular to these masks, ensuring not to intersect with neighboring cells. Line profiles that intersected with the Hoechst fluorescence in the nuclei were automatically excluded from analysis. Manual verification of the NG cell masks and line profile positions was performed using a custom graphical interface developed with Napari. Only cells and line profiles that passed the manual verification were included into the subsequent analysis. Cells with fewer than three confirmed line profiles were automatically excluded. Fluorescence data from the EYFP channel were recorded along each line profile and averaged across all profiles for each cell, resulting in an expression line profile for each cell. These expression line profiles from all verified cells were then averaged, and standard deviations were calculated.

In order to avoid averaging of the fluorescence from multiple cells only sectors of each cell (centered on the cell’s centroid) that were not oriented towards adjacent cells were selected for analysis. The average fluorescence in the EYFP channel within these sectors was subsequently used to calculate the corrected partial cell fluorescence per area (CPCF per area). Only sectors with an angle greater than 120 degrees were included in the analysis. The CPCF per area value is a measure for the average cell fluorescence and represents an approximation of the overall ChR expression.

#### Supplementary Note 12: rAAV purification and cloning of constructs

A triple transfection of HEK-293T cells was performed using the pHelper plasmid (TaKaRa, USA), trans-plasmid with either the AAV2/9, PHP.eB or PHP.S capsid[56] and cis-plasmid with either ChRmine or ChReef under the control of human synapsin or CAG promotor. Cells were regularly tested for mycoplasma contamination. Viral particles were harvested cells and medium; precipitated from medium with 40% polyethylene glycol 8000 (Acros Organics, Germany) in 500mM NaCl and combined with cell pellets for processing. Pellets were suspended in 500mM NaCl, 40mM Tris, 2.5mM MgCl2, pH 8, and 100 Uml−1 of salt-activated nuclease (Arcticzymes, USA) at 37 °C for 30 min. Cleared lysates were purified over iodixanol (Optiprep, Axis Shield, Norway) step gradients (15%, 25%, 40%, and 60%) at 350,000 × g for 2.25 h. rAAV containing fractions were concentrated using Amicon filters (EMD, UFC910024) and formulated in sterile phosphate-buffered saline (PBS) supplemented with 0.001% Pluronic F-68 (Gibco, Germany). Viral vector titers were determined by the number of DNase I resistant vg using AAV titration kit (TaKaRa/Clontech) by qPCR (StepOne, Applied Biosystems) according the manufacturer's instructions or with adaptations to usage in crystal digital PCR (dPCR) system. Here, dilutions of purified vector DNA were subjected to crystal dPCR using 5x Naica PCR reaction mix (Stilla, R10056) supplemented with primers provided in the AAV titration kit, 0,8mg/µl Alexa 647 (Invitrogen, 11570266) and 1.5x Eva Green (Biotum, #31000-T). Formation of droplet crystals and PCR was performed in a Naica Geode (Stilla) using Naica Ruby or Sapphire Chips (Stilla) and finally analyzed in the Naica Prism3 reader (Stilla) equipped with Crystal Reader and Crystal Miner softwares (Stilla). rAAVs titers and quantification method used is indicated for each rAAV in the respective methods chapter. Purity of produced viral vectors was routinely checked by silver staining (Pierce, Germany) after gel electrophoresis (Novex™ 4–12% Tris–Glycine, Thermo Fisher Scientific) according to the manufacturer's instruction. Viral vector stocks were kept at −80 °C until injection.

#### Supplementary Note 13: Preparing neonatal mouse cardiomyocytes

Hearts were explanted from p0-p3 mice killed by decapitation and directly placed on ice cold PBS. Remaining tissue and atria were carefully removed, and ventricular CMs isolated with the neonatal heart dissociation kit for mouse and rat (Miltenyi, Germany). 1.5 – 2 million cells were plated on a 6-well-plate coated with 10 µg/ml fibronectin (Sigma; F1141-5MG) and maintained in IMDM (GlutaMAX, no phenol red; Gibco 21056-023) supplemented with 10% FBS superior (Sigma; S0615) and 1% Penicillin/Streptomycin (Life Technologies; 15140122) in standard cultivation conditions (37°C, 5% $CO_2$). One day later, medium was exchanged to reduce FBS to 0.5% and AAV transduction was performed with AAV2/9-CAG-ChRmine-TS-EYFP-ES with $1.1 \times 10^{10}$ GC/ml (qPCR) and AAV2/9-CAG-ChReef-TS-EYFP-ES with $6 \times 10^{10}$ GC/ml (qPCR) corresponding to ~226 and ~69 vp/cell, respectively. On day 3, the cells were dissociated and plated on Ø 10 mm round coverslips in CM clusters compromising 50,000 cells within 15 µl locally on an 8 µl 0.1% gelatine drop. Cells were incubated in this small drop for 20 min to build cardiac clusters and afterwards medium filled up in the well. After 1-2 days CM cluster started to beat spontaneously and 0.5% FBS IMDM medium was exchanged every 2-3 days.

#### Supplementary Note 14: Preparing hiPSC-derived cardiomyocytes

Human CMs were differentiated from the induced pluripotent stem cell line UMGi014-A clone 2, which was created from peripheral mononuclear blood cells from a healthy male donor using integration-free Sendai virus and was described previously. CMs differentiation and purification involved WNT signaling modulation and lactate-based metabolic selection as described previously[57,58]. Differentiated CMs were cryopreserved for further use. For measurements, 1.5 – 2 million cells were thawed and plated on a 6-well-plate coated with 16 mg/mL Geltrex (Thermo Fisher Scientific; A1413302) and maintained in CM culture medium

(RPMI 1640 medium (GlutaMAX; Thermo Fisher Scientific; 72400054), B27 (Thermo Fisher Scientific; 1750404) supplemented with 2 mM thiazovivin (Sigma-Aldrich; 420220-10MG), at 37°C and 5% $CO_2$. Medium was replaced after two days to remove residues of thiazovivin and changed every 2-3 days afterwards. AAV transduction was performed with MyoAAV-CAG-ChRmine-TS-EYFP-ES with ~$1x10^{11}$ GC or MyoAAV-CAG-ChReef-TS-EYFP-ES with ~$1.8x10^{10}$ GC (CdPCR). Four days after transduction, cells were dissociated and plated on Ø10 mm round coverslips in cardiac cell clusters comprising 50,000 cells within 8 µl drop on a Geltrex/DMEM/Ham's F-12 coated area. Cells were incubated in this drop for 40 min to build CM clusters and afterwards medium was filled up with CM culture medium. After two days CM clusters started to beat spontaneously. Medium was replaced every 2-3 days.

**Supplementary Note 15: Histology for cardiomyocytes**

Neonatal heart cells as well as hiPSC-derived CMs were fixed in 4% formaldehyde for 20 min, permeabilised with 0.2% Triton X100 for 20 min and stained in DPBS (Sigma; D8537) supplemented with 5% donkey serum for 2 h with primary antibodies against cardiac troponin I (ab47003, Abcam: 1:800) and for 1 h with secondary antibodies conjugated with Cy5 (711-175-152, JacksonLab, U.S., 1:400) diluted in DPBS with 1:1000 DAPI (0018860.01, Th.Geyer) at room temperature. Images of single CMs for analysis of expression rate were taken through a 10x (10X/NA 0,4 Olympus UPLSAPO) or 20x objective (20x/NA 0,7 Olympus UCPLFLN PH) with an IX83 inverted fluorescence microscope equipped with an ORCA-flash 4.0 digital camera (C11440, Hamamatsu Photonics) and the MT20 illumination system as light source controlled via the CellSens® software. Acquisition of images was performed with following filter settings: 387/11 excitation, 410 beamsplitter and 440/40 emission for DAPI, 485/20, 504 and 525/30 for eYFP, 560/25, 582, and 684/24 for Cy5. Transduction rate was calculated by the percentage of eYFP positive cells among all CMs identified by cardiac troponin I antibody-staining.

CM clusters were imaged with a LSM 800 confocal microscope (Zeiss) equipped with spectral multi-alkali photomultiplier detectors. Z-stack acquisition was performed using a Plan-Apochromat 20x/0.8 M27 via the ZEN 2.6 (blue edition) software (Zeiss) with a pinhole of 55 µm for Dapi, 24 µm for eYFP, 27 µm for the red channel and 31 µm to image Cy5. Pictures were post-processed using stack correction (Background+Decay+Flickr), extended depth of focus (Wavelets, highest Z-Stack Alignment) and Image calculator (subtraction of autofluorescence) of the Zen 3.4 (blue edition) software.

**Supplementary Note 16: Intravitreal injection**

All *in vivo* electrophysiological experiments were carried out in compliance with the relevant national and international guidelines (European Guideline for animal experiments 2010/63/EU) as well as in accordance with German laws governing animal use. The procedures have been approved by the responsible regional government office. Rodents were kept in a 12h light/dark cycle with ad libitum access to food and water.

3-months old female and male Rd1 (C3HeB/FeJ, JAX 000658) mice were anesthetized with a subcutaneous injection of a fentanyl (0.05 mg/kg), midazolam (5 mg/kg), and medetomidine (0.5 mg/kg) (FMM) cocktail. Mice were then secured using a bite bar and placed on top of a temperature controller (Supertech) to maintain a body temperature of 37°C. For local anesthesia of the eyes, eye drops containing 0.5% Proxymetacain were applied and hydration gel (Bayer, Bepanthen) was placed on the eyes to prevent dryness whenever possible. Before injection, a hole posterior to the ora serrata was created using a 27-gauge needle. Then, a Hamilton syringe was guided through this hole into the intravitreal space and 2µl of virus solution of PHP.eB-hsyn-ChRmine-TS-EYFP-ES (1,89x$10^{13}$ GC/ml, dPCR) or PHP.eB-hsyn-ChReef-TS-EYFP-ES (8,59x$10^{13}$ GC/ml, dPCR) was injected at 4µl/min using a motorized injector (Stoelting). After 5 min the syringe was withdrawn and the anesthesia was reversed with a subcutaneous injection of a naloxone (1.2 mg/kg), flumazenil (0.5 mg/kg), and atipamezole (2.5 mg/kg) cocktail.

**Supplementary Note 17: Analysis of neural responses to optogenetic stimulation of the visual system**

Only units with stable responses across the duration of the session and high kilosort quality scores were included in subsequent analysis. To obtain continuous firing rate estimates, single unit responses were binned with a bin size of 1 ms and convolved with a gaussian kernel with a standard deviation of 10 ms. Trial-averaged responses were normalized to the mean and standard deviation of the baseline (400ms before stimulus onset) across all intensities. Units that did not respond to the highest light intensity were excluded (mean normalized firing rate during the stimulus smaller than 2.58). The Gramm toolbox[62] was used to visualize raster plots. For significance testing across time points a permutation based method[63] was used as described[64]. Briefly, using unpaired t-tests differences between the two conditions for each time point are determined and subsequently time clusters are identified using the bwconncomp.m function in Matlab (4 connected components). Time cluster significance is determined by comparison the t-statistics of the actual data against the t-statistics of shuffled data (10000 permutations with randomized labels).

**Supplementary Note 18: Histology and imaging after optogenetic stimulation of the visual system**

After washing in PBS, retinas were prepared and fixed again for 10 min in 4% PFA at RT. Fixed and washed retinas were put in 30% Sucrose and subjected two three rounds of freezing and thawing on dry ice. Then retinas were incubated for 1h in 10% normal donkey serum (NDS) and subsequently in primary antibody solution (rabbit-GFP (Proteintech, 50430-2-AP), 1:666 in PBS with 3% NDS and 0.5% Triton-X) for 4 days at 4°C. After incubation, retinas were washed in PBS and incubated in secondary antibody overnight at 4°C (AlexaFluor 488 (Invitrogen, A32970), 1:500 in PBS with 3% NDS and 0.5% Triton-X). Finally, washed retinas were mounted in DAPI-containing Vectashield (Vector Laboratories, H-1200).

For post-hoc analysis of recording sites, brains were collected and incubated in 4% PFA at 4°C overnight. Then brains were washed and embedded in 4% agarose. 100µm coronal slices were cut using a vibratome (Leica microsystems, VT1000S) and stained for 2h at RT (DAPI, Thermo Fisher Scientific, D1306, 2.5 µg/ml). Washed brain slices were mounted on microscope slides in Vectashield (Vector Laboratories, H-1200).

Images were captured using a Leica TCS SP8 laser scanning confocal microscope with 63×/ 20x oil immersion or a 10× air objectives (Leica Microsystems). Images were merged and processed using LAS X (Leica Microsystems) and Adobe Photoshop software.

**Supplementary Note 19: Behaviour experiments for vision restoration in mice**

Mice of both sexes were used and experiments were performed during the daytime (corresponding to the mouse's subjective nighttime). All procedures related to the behaviour experiments were performed in accordance with the Canadian Council on Animal Care and approved by the Montreal Neurological Institute's Animal Care Committee.

**Supplementary Note 20: Postnatal AAV injection into the cochlea in mice and Mongolian gerbils**

The same injection approach was performed for all animals which later would be subject to either: auditory brainstem recordings, juxtacellular recordings from single putative SGNs, *ex vivo* acute slice electrophysiology in mice, or recordings from the inferior colliculus in Mongolian gerbils. AAV injections into the left ear were performed at postnatal day 6 in C57BL/6J wild-type or B6;129P2-Otof$^{tm1.1Erei}$ (MGI: 4950055 [66]) mice or at postnatal day 7.3 ± 1.3 days in Mongolian gerbils[18,55]. The right ear served as a non-injected control. In brief, mouse and gerbil pups were randomly selected for virus injections. Under general isoflurane anesthesia (5% for anesthesia induction, 1–3% for maintenance with frequent monitoring of the hind-limb withdrawal reflex and anesthesia adjustments, accordingly) and local xylocaine as well as buprenorphine (0.1 mg/kg) and carprofen (5 mg/kg) administration for analgesia, the cochlea of the left ear was accessed via a retroauricular incision. Body temperature was maintained at physiological temperature. The cochlea was gently punctured using a quartz capillary pipette, which was kept in place to inject the virus of following titers:

Mice: AAV2/9-hsyn-ChRmine-TS-EYFP-ES 5.1x$10^{13}$ GC/ml (qPCR),
AAV2/9-hsyn-ChReef-TS-EYFP-ES 3.1x$10^{13}$ (qPCR) and 2.45x$10^{13}$ GC/ml (dPCR)

Gerbil: PHP.S-hsyn-ChReef-TS-EYFP-ES 5.25 x$10^{13}$GC/ml (qPCR)

After virus injection the tissue in the injection area was repositioned and the wound was sutured. Carprofen (5 mg/kg) was administered for analgesia the day after surgery.

**Supplementary Note 21: Description of anesthesia and analgesia for adult rodents undergoing *in vivo* optical stimulation**

Surgeries and measurements were performed under anesthesia using isoflurane (5% for anesthesia induction, 1–3% for maintenance with frequent monitoring of the hind-limb withdrawal reflex and anesthesia adjustments, accordingly) and analgesia by sub-cutaneous injections of buprenorphine (0.1 mg/kg body weight) and meloxicam (5 mg/kg body weight). Body temperature was maintained at 37°C using a custom-designed heat plate through all the procedures.

**Supplementary Note 22: Optical stimulation of the auditory pathway in rodents *in vivo***

The left cochlea was exposed by performing a retroauricular incision behind the pinna followed by a bullostomy, where the round window was visualized and punctured. For mice a 50 µm optical fiber coupled to a 594 nm (OBIS LS OPSL, 100 mW, Coherent Inc., Santa Clara, United States) or a 522 nm (Oxxius combined with a motorized Power Attenuator, Lannion, France) diode laser was used. For gerbils a 200 µm optical fiber was only coupled to the prior mentioned 522 nm laser. Laser power was calibrated prior to each experiment using a laser power meter (Thorlabs PM100USB).

**Supplementary Note 23: Kanamycin deafness model in mice and Mongolian gerbil**

For the kanamycin deafness model, the cochlea was deafened by delivering 200 mg/ml of kanamycin through the round window via a microfile needle, acutely. It was left for approximately 7 minutes (for mice) or 15 to 30 minutes (for gerbils) before additional measurements were performed to confirm deafness by absence of acoustically evoked auditory brainstem responses. The post and pre-deafened auditory brainstem responses were compared to evaluate the effectiveness of the deafening procedure.

**Supplementary Note 24: Experimental set-up for optical *in vivo* recordings in rodents**

Stimulus generation and delivery, as well as data acquisition was performed using a custom-written software (MATLAB, MathWorks, Natick, MA, United States) employing National Instrument data acquisition cards and a custom-build laser-controller. Recordings were conducted in a soundproof chamber (IAC Acoustics, IL, United States).

**Supplementary Note 25: Data analysis of acoustic and optically evoked auditory brainstem responses in mice**

The mouse ABR data was analyzed using custom-made Matlab software (The MathWorks, Inc., Natick, MA, USA). Averages were expressed as mean ± standard deviation. For statistical comparison between two groups, data sets were tested for equality of variances (F-test) followed by two-tailed unpaired Student’s t-test.

#### Supplementary Note 26: Procedures for immunostaining cochleae of mice and Mongolian gerbils

At the end of all *in vivo* recordings of rodents, the animals were sacrificed under deep anesthesia via cervical dislocation. The cochleae were collected and fixed in 4% formaldehyde solution for 45 minutes. They were then decalcified for 3-4 days in 0.12 M ethylenediaminetetraacetic acid (EDTA), dehydrated in 25% sucrose solution for 24 hours and cryosectioned (mid-modiolar cryosections, 25 µm tick). Immunofluorescence staining were done in 16% goat serum dilution buffer (16% normal goat serum, 450 mM NaCl, 0.6% Triton X-100, 20 mM phosphate buffer, pH 7.4). The following primary antibodies were used at 4 °C overnight: chicken anti-GFP (1:500, ab13970 Abcam, USA) and guinea pig anti-parvalbumin (1:300, 195004 Synaptic Systems, Germany); and secondary antibodies for one hour in room temperature: goat anti-chicken 488 IgG (1:200, A-11039 Thermo Fisher Scientific, USA) and goat anti-guinea pig 568 IgG (1:200, A-1107 Thermo Fisher Scientific, USA). Finally, slices were mounted in Mowiol 4-88 (Carl Roth, Germany). Samples were imaged with a confocal microscope SP8 (Leica, Germany) mounted with a 40x oil objective. For each cochlear turn (apex, middle and base) an image was taken focusing on the modiolus.

#### Supplementary Note 27: Preparation of brain stem slices

The brain was immediately immersed in ice cold cutting solution (Supplementary Table 8) and pinned, dorsal side down, on a wax stage. With the ventral side of the brainstem exposed, the meninges were removed with curved forceps and a mid-sagittal cut was performed using a straight razor, splitting the brain in two hemispheres. Scissors were used to cut between the cerebellum and the occipital lobe of each hemisphere in order to separate the hind-brain from the fore-brain. The two blocks of hind-brain, containing brain-stem and cerebellum, were glued with cyanoacrylate glue, (Loctite 401, Henkel) medial side down, on the stage of a Leica (Wetzlar, Germany) VT 1200 vibratome. The lateral side of both brain blocks was facing upwards and was facing towards the advancing blade. A vibration amplitude of 1.5mm was used and the blade was placed at the height of the cerebellar flocculus. Cerebellar tissue was sectioned at an advancing speed of 0.05 – 0.1 mm/s and discarded. Thereafter, a speed of 0.02mm/s was used to obtain 150µm thick parasagittal slices of the cochlear nucleus. The slices were incubated in artificial cerebro-spinal fluid (aCSF, Table S8) for a total of 30 minutes in a 35°C water bath (Julabo, Seelbach, Germany), after which they were maintained in aCSF at room temperature (23 – 25 °C) until recording. The pH of both the cutting solution and the aCSF was adjusted to 7.4. The osmolarity was, in mOsm, 310 for the aCSF and 320 for the cutting solution. Both solutions were continuously aerated with carbogen (95% $O_2$, 5% $CO_2$).

**Supplementary Note 28: Auditory brainstem slice electrophysiology**

The sampling interval was set to 25µs and the filter at 7.3 kHz. For better visualization of the slice's cell layers and auditory nerve fibers (ANFs), differential interference contrast (DIC) microscopy was used on upright Olympus BX51WI microscope with a water immersion LUMPLFLN40XW objective (Olympus, Hamburg, Germany). All recordings were performed at near physiological temperature (33 – 35°C). The bath solution flowed through an inline solution heater (SC-20 with CL-200A controller; Warner Instruments, Hamden, CT, USA). The temperature of the solution was monitored by a thermistor placed between the opening of the inflow tube and the slice in the recording chamber.

Borosilicate filamentous glass capillaries (GB150F, inner diameter: 0.86 mm, outer diameter: 1.50 mm, length: 80 mm, Science Products GmbH, Hofheim, Germany) were used to pull patch pipettes on a P-1000 micropipette puller (Sutter Instruments Co., Novato, CA, USA). The resistance at the tip of the pipette was strictly 2 – 3 MΩ to avoid increases of the series resistance ($R_s$) over long recordings. The pipettes were filled with intracellular solution (details on Table S8), which additionally contained 1 mM of Alexa-568 (Invitrogen / Fisher Scientific, Schwerte, Germany). The pH of the solution was adjusted to 7.4 and its osmolarity to 325 mOsm, to prevent osmotic collapse of the patched neuron, preserve the cell's vigour and prevent fluctuations of the $R_s$ over long recordings. Voltage clamp experiments were performed with a $V_{hold}$ = -70 mV, corrected for a liquid junction potential of 12 mV. Only cells with $R_s$ values less than 8 MΩ were included in the analysis. Afferent fiber optogenetic stimulation was performed using a green laser included in the Oxxius multi-wavelength bundle (LBX-488-40-CSB, maximum Power: 40 mW; LCX-532-50CSB, maximum power: 50 mW; OBIS-594-100, maximum power: 100 mW; LBX-638-100-CSB, maximum power: 100mW; Laser 2000 GmbH, Wessling, Germany). Synapses were driven with trains of stimuli yielding varying success rates of optically eliciting excitatory postsynaptic currents, hereafter termed oEPSC probability ($P_{oEPSC}$). $P_{oEPSC}$ was defined as $\frac{Number\ of\ oEPSCS}{50}$, 50 signifying the number if 1ms light pulses that were delivered to the ANF. Minimal stimulation irradiance was defined as the lowest value that could yield a $P_{oEPSC}$ = 1 at 10 Hz. This irradiance value was used later in the experiment to assess the $P_{oEPSC}$ at higher frequencies. For ANF stimulation, 5, 10, 20, 33, 40, 50 and 100 Hz were the tested frequencies. In direct transduction experiments, where the photocurrent properties of ChReef were measured, 10, 20, 30, 50, 100, 200, 300 and 350 Hz were used instead. Electrophysiology data were analysed using Igor Pro (Wavemetrics, Lake Oswego, OR, USA). Evoked release was analysed using custom Igor scripts. Confocal z-stacks were processed with the NIH ImageJ software[68]. Figures were assembled for display using open-source vector graphic software Inkscape (https://inkscape.org).

**Supplementary Note 29: Immunohistochemistry and immunofluorescence imaging for acute slice electrophysiology**

Afterwards they were washed in PBS for 10 minutes to halt fixation and blocked with GSDB for 1 hour at room temperature. After fixation the slides were kept for 10 minutes in ice cold PBS to halt fixation and then washed for 1 hour at room temperature in goat serum dilution buffer, which contained in 16% normal goat serum, 450 mM NaCl, 0.3% Triton X-100, and 20 mM phosphate buffer (PB, pH 7.4). The slides were then incubated in primary antibodies diluted in GSDB, for 3 hours, in a wet chamber at room temperature. Four washing steps followed, that lasted 10 minutes each, two of them with Wash Buffer (450 mM NaCl, 0.3% Triton X-100 and 20mM PB) and two more with PBS. The slides were then incubated with secondary antibodies overnight at 4°C and, afterwards, washed twice with Wash Buffer and twice more with PBS as before. Before mounting, the slides were maintained in 5mM PB for 10 minutes and were then mounted with a drop of Mowiol (Carl Roth GmbH, Karlsruhe, Germany) and covered with a thin glass coverslip. Double sided tape spacers of 120μm thickness were used for mounting to flank the slice before the application of Mowiol. This way, the coverslip's weight would not be applied fully on the slice, avoiding extra vertical pressure, which resulted in better preservation of structural integrity of the tissue. Immunofluorescence imaging was performed on a Zeiss LSM 780 Microscope with a 40x NA1.4 *Oil* Plan-Apochromat objective. *Primary antibodies:* chicken-anti-GFP (1:500, Abcam, Berlin, Germany), guinea-pig-anti-vGLUT1 (1:1000, Synaptic Systems GmbH, Göttingen, Germany). *Secondary antibodies:* goat-anti-chicken 488 (1:200, Thermo Fisher Scientific, Waltham, USA) goat-anti-guinea-pig 568 (1:200, Thermo Fisher Scientific, Waltham, USA).

**Supplementary Note 30: Cumulative discrimination index (d') calculation**

Post-stimulus time histograms (PSTH) were computed by binning spike counts in 0.2 ms bins. The first three consecutive bins crossing threshold of 3x MAD indicate $t_{on}$ and, respectively, the last three consecutive bins over threshold indicate $t_{off}$. Based on the time window extracted, the signal was further analyzed and sorted in a two-dimensional matrix according to stimulus intensity and corresponding recording site. From this matrix, we calculated the cumulative distribution index (d') based on spike rates calculated across increasing intensities starting with 0 intensity condition.

**Supplementary Note 31: Temporal analysis of neural clusters**

To calculate the number of spikes per stimulus, we divided the spike counts recorded by the number of repeats per recording and by the number of pulses from stimulation rate used. See supp,ecalculate the vector strength, a measure of spike timing relative to stimulation cycle the following formula was previously[40]:

$$VS = \frac{\sqrt{[\sum_{i=1}^{n} \cos \theta i]^2 + [\sum_{i=1}^{n} \sin \theta i]^2}}{n}$$ [Equation 7]

With θ describing the spike timing computed from stimulus onset to following stimulus onset for a given stimulus cycle (phase). The Rayleigh test (eq. 8) was then performed in order to test the significance of vector strength for each unit and each repeat.

$$L = 2 * length(\theta i) * (VS^2)$$ [Equation 8]

If L<13.8, p>0.001 and vector strength significance is rejected and set to 0. Firing rates were computed across all recordings by dividing the spike counts recorded in the determined spike window by the duration of that spike window. These values were normalized to the highest firing rate recorded for plotting.

**Supplementary Note 32: Subcellular expression profiles for SGNs in cochlear cryosections**

To assess the subcellular expression profiles of ChRmine and ChReef in SGNs, we used a custom-written Python program to plot fluorescence line profiles, incorporating a manual verification process. The process began by using the manually detected centroids (using custom MATLAB script as explained in methods section) of all cells to generate 2D square crops of the images, each containing one complete SGN. For each 2D crop, the Z-plane was selected by maximizing fluorescence in the cytoplasmic parvalbumin channel.

Cell membrane masks were automatically generated based on the cytoplasmic parvalbumin channel fluorescence using a fixed threshold. Line profiles were then automatically positioned perpendicular to these masks and avoiding intersection with surrounding cells. Manual verification of SGN masks and line profile positions was conducted using a custom graphical interface developed with Napari. The Z-plane was manually checked for a noticeable fluorescence drop indicating the position of the nucleus; cells lacking a discernible nucleus were excluded to prevent bias from including line profiles in the nuclear region or parallel to the cell surface. Any line profiles that intersected visually identifiable nuclei were excluded from further analysis.

Only cells and line profiles that passed manual verification were used for further analysis. Cells with fewer than three confirmed line profiles were automatically excluded. Fluorescence data from the YFP channel was recorded along each line profile and averaged across all lines for each cell, creating an expression line profile for each cell. These expression line profiles from all verified cells were then averaged, and the standard deviation was calculated.

**Supplementary Note 33: Brain histology**

Brains were left in 8% FA over a week and transferred to 30% sucrose for 24 hours. Brains were then carefully mounted using cryomatrix gel (Shandon Cryomatrix, Thermo Fisher Scientific, Waltham MA, USA) and sectioned coronally. Slices were 50 µm thick, transferred onto microscope slides (Thermo Superfrost/ColorFrost Plus, Charged 90° - CS/1440CS/1440, Erie Scientific, Ramsey MI, USA) and prepared for antibody staining. The staining follows the cochleae staining protocol, after PBS washing, sections were blocked with goat serum dilution buffer (GSDB) for 1h at room temperature before primary antibodies for parvalbumin (1:300, guinea pig, Synaptic Systems, Goettingen, Germany) and GFP (to label ChReef, 1:500, chicken, Abcam, Cambridge, UK). Goat anti-guinea-pig-568 and goat anti-chicken-488 were used for secondary antibodies and incubated at room temperature for 1h – slides were ultimately mounted with Mowiol and imaged the following day. Images were taken in oil at 40x using LSM510 (Zeiss, 467 Jena, Germany) microscope and analyzed with ImageJ (U. S. National Institutes of Health, Bethesda, Maryland, USA).

##### Supplementary Note 34: Animal housing

Data were obtained from nine 2.5 to 3.75 year old common marmosets of either sex (5 male, 4 female). Animals were pair housed in a colony of about 50 animals consisting of several rooms with acoustic but no visual contact to neighboring pairs. The colony maintained a 12h:12h light-dark cycle at a temperature of 26 ± 1.5 °C and a relative humidity of 60 - 80 %. Animals were fed *ad libitum* a diet of fresh fruits, vegetables, nuts, arabic gum, rice, pasta, potatoes, meal worms, crickets, joghurt, various minerals as well as pellets. Water was always available and was occasionally mixed up with unsweetened tea.

##### Supplementary Note 35: Acoustic auditory brainstem recordings

Animals were anesthetized with a mixture of Alfaxan (10mg/kg BW, i.m.) and Midazolam (0.125 mg per animal, i.m.). Further, Metacam (0.2 mg/kg BW) was administered either orally – or if not possible – via s.c. injection. After anesthesia initiation, animals were placed on a heat pad, wrapped in a forced air warming blanket and body temperature was maintained at 37 to 38 °C. Otoscopic evaluation of ear canals was performed next to rule out signs of infection and was found to be normal in all cases. Afterwards, animals were transferred onto a warmed saline bag on a heat pad within a stereotaxic frame. Animals were not fixed inside the frame but made use of custom-made ear bars containing earphone (Etymotic ER-4PT) and microphone (Sennheiser MKE 2 P-C) inserts. Ear bars were gently inserted into the outer ear canals. Monitoring of the microphone signal upon acoustic stimulation ensured correct placement. Auditory stimuli were calibrated separately with 3D printed ear molds of common marmosets where a 1/8$^{th}$ inch microphone (G.R.A.S. 46 BE) was placed at the position of the tympanic membrane. Stimulus generation and delivery, as well as data acquisition was performed using a custom-written software (MATLAB, MathWorks, Natick, MA, United States) employing National Instrument data acquisition cards and a custom-build laser-controller. Recordings were conducted in a dedicated operating theater for non-human primate surgery. If necessary, anesthesia was maintained by re-injecting Alfaxan (5 mg/kg BW). Acoustically evoked ABRs (aABRs) were recorded by placing needle electrodes behind the pinna, on the vertex, and on the neck of the anesthetized animals. The difference in potential between the vertex and mastoid subdermal needles was amplified using a custom-designed amplifier, sampled at a rate of 50 kHz for 10 ms, filtered (300–3000 Hz) and averaged across at least 750 stimulus presentations. In case of pure tone stimulation a bandpass filter centered on the pure tone frequency removed stimulus artifacts observed occasionally. The ABRs threshold was defined and determined as the lowest light or sound intensity for which one of the 5 waves (aABR) or 2 waves (oABR) was reliably visible.

#### Supplementary Note 36: Injection method

First, animals received Metacam (0.2 mg/kg BW) either orally – or if not possible – via s.c. injection and were anesthetized with injections of Alfaxan (10mg/kg BW, i.m.) and Midazolam (0.125 mg per animal, i.m.). Additionally, dexamethasone (5 mg/kg BW, i.m.) and Cerenia (1 mg/kg BW, s.c.) were also applied. The fur around the surgical site was removed. Then, an intravenous catheter was placed in the vena saphena and the animal was intubated using a 00 Miller blade and a custom-made unblocked endotracheal tube for artificial ventilation. Next, anesthesia was transferred to a combined volatile-intravenous anesthesia with Sevoflurane (ca. 2 – 3 %) and Propofol-Remifentanil (0.3 – 0.4 mg/kg/min; 10 – 25 µg/kg/h, respectively) to effect. During surgery, body temperature was maintained between 37 and 38°C with a heating pad and a forced air warming blanket wrapped around the animals' body. Additionally, physiological parameters were monitored throughout the surgery via ECG and pulse oximetry. Surgery itself was performed under standard aseptic technique. Local analgesia was achieved with Carbostesin injections around the surgical site. The cochlea was accessed via a retroauricular incision, blunt dissection of post-auricular as well as temporal muscles to expose the linea temporalis and mastoidectomy ventral to the linea temporalis. Utmost care was used to avoid damaging the horizontal semicircular canal and facial nerve which served as guiding landmarks. Animals were initially deafened with flushes of 10 % Neomycin as described previously [60]. In later experiments, Neomycin was added to the viral suspension to a final concentration of 10 %. In addition, the ossicle chain was disrupted by removal of the malleus. The viral suspension was administered either via a pipette pulled from borosilicate and pressure injection or a silicone catheter was used to slowly (15 µl/30 min; UMP3, World Precision Instruments) inject. In both cases, the round window membrane was identified and gently ruptured. To prevent pressure build-up inside the cochlea during slow injection, a ventilation hole was opened either with a rose burr next to the oval window or by removing the stapes when surgical access was favorable. Excess viral suspension was carefully suctioned to avoid unwanted transfection of surrounding tissue. In later experiments, Dexamethasone was added to the viral suspension (2 %). Three different titers of viral suspension (15µl) were used 5x $10^{10}$ , $10^{11}$, und 5x $10^{11}$ GC) and injected in three animals each. Animals were randomly assigned to each group and experimenters were blinded to the condition. After virus injection, the round window and vent holes were closed with autologous tissue, the mastoidectomy closed by replacing the temporal and post-auricular muscles. The skin and muscles were then sutured with 5/0 absorbable suture. For direct post-operative pain management, Buprenorphine (0.005 mg/kg BW, s.c.) was administered. To prevent hypoglycemia Glucose (5%, 2 ml) was administered s.c. Post-operative care included repeated antibiotic administration (Amoxicillin 37.5 mg/kg BW) every 48 h for 8 days as well as analgesia via Metacam administration (0.2 mkg/kg BW, p.o.).

**Supplementary Note 37: Immunostaining and confocal imaging of cochlear cryosections**

The facial nerves exiting the foramen stylomastoideus were dissected and the soft tissue overlying the temporal bones were removed to allow trimming bone around the cochlea under visual guidance. Next, the cochlea and vestibular organs were transferred into 5 M EDTA for decalcification. During a period of at least 2 weeks, EDTA was regularly exchanged and softened bone removed from the cochlea. Additionally, the facial nerve was removed from its bony capsule overlying the cochlea. Once completed, the cochlea was prepared for whole mounts of the organ of Corti and mid-modiolar cochlear cryosections. Here, the spiral ligament and bone lamella of the apex were carefully removed to access the organ of Corti and dissecting it by cutting the spiral lamina. Next, the stria vascularis and tectorial membrane were removed. This procedure was repeated until the whole organ of Corti was dissected into consecutive whole mounts and the cochlear modiolus. Finally, the whole mounts and modioli were stored in phosphate buffered saline. The modioli were then processed similarly to rodent samples: after a 24 h dehydration step in 25 % sucrose solution, samples were cryosectioned (mid-modiolar cryosections, 16 µm tick). Immunofluorescence staining was performed in 16% goat serum dilution buffer (16% normal goat serum, 450 mM NaCl, 0.6% Triton X-100, 20 mM phosphate buffer, pH 7.4). The following primary antibodies were used at 4 °C overnight: chicken anti-GFP (1:500, ab13970 Abcam, USA) and guinea pig anti-parvalbumin (1:300, 195004 Synaptic Systems, Germany) as well as mouse anti-NF200 (1:400, Sigma, St. Louis, USA); and secondary antibodies for one hour in room temperature: goat anti-chicken 488 IgG (1:200, Invitrogen Scientific, USA), goat anti-guinea pig 568 IgG (1:200, Invitrogen, USA) and anti-mouse 633 (1:200, Invitrogen). Finally, slices were mounted in Mowiol 4-88 (Carl Roth, Germany). Whole mounts were first permeabilized with TritonX 0,5% in PBS. Staining was then performed in 10 % goat serum dilution buffer and guinea pig anti-parvalbumin (1:200; Synaptic systems, Germany), rabbit anti-Otoferlin (1:500; SySy) and chicken anti-GFP (1:500; Abcam) were used as primary antibodies. As secondary antibodies anti-chicken 488 (1:1000; Invitrogen), anti-guinea pig 568 (1:1000; Invitrogen) and anti-rabbit 633 (1:1000; Thermo Fisher) were used. Samples were imaged with a confocal microscope SP8 (Leica, Germany) mounted with a 40x oil objective. For each cochlear turn (apex, middle and base) an image was taken focusing on the modiolus. Image analysis was performed by a custom-written MATLAB script modified from[61]. Briefly, SGN somas and modiolus area were manually detected using a touch screen from the parvalbumin images. Next, individual somas were automatically segmented using Otsu's threshold method from every Z-stack and a mask corresponding to the given SGN was defined for the Z-stack for which the mask was fulfilling the criteria of size (area and diameter) and circularity. In case the segmentation was not deemed to be sufficient, the segmentation of the given SGN soma was performed manually. Next, the median GFP brightness of each SGN was measured and its distribution was fitted with a Gaussian mixture model with up to 3 components. A threshold, above which SGNs somas were considered as transduced, was defined as average + 3 x standard deviation of the Gaussian distribution with the lowest mean. The density of inner hair cells (IHCs) along the whole mounts was calculated by placing a curve along the (presumed) IHC, measuring its length and assessing the number of identified IHC.
